# Targeting UBE3A and downstream estrogen receptor-β signaling to restore oligodendroglial homeostasis in Angelman syndrome

**DOI:** 10.64898/2026.05.21.726878

**Authors:** Xin Yang, Sonia Mayoral, Yong-Hui Jiang, John Marshall, Yu-Wen Alvin Huang

## Abstract

Mutations that reduce UBE3A cause Angelman syndrome (AS), a neurodevelopmental disorder marked by severe developmental delay and neuropsychiatric symptoms. Although UBE3A has been studied primarily in neurons, it is also expressed in the oligodendrocyte lineage, raising the possibility that glial dysfunction contributes to disease phenotypes. Here we identify an intrinsic, UBE3A-dependent mechanism that governs oligodendrocyte precursor cell (OPC) homeostasis, a process necessary for normal myelination. Using human iPSC-derived OPCs, we performed a targeted compound screen and found that loss of UBE3A diminishes estrogen receptor-β (ESRβ) signaling, leading to impaired OPC self-renewal. Mechanistically, UBE3A sustains ESRβ levels in OPCs and thereby maintains downstream signaling required for OPC proliferation and stable OPC density. To address cell-type specificity and neuron-glia interactions, we combined oligodendrocyte-lineage-restricted *Ube3a* knockdown *in vivo* with reductionist iPSC neuron-OPC co-culture assays, which together support an intrinsic requirement for oligodendroglial UBE3A in OPC proliferation and myelination. Pharmacologic activation of ESRβ with selective agonists restored OPC self-renewal and myelination deficits in AS patient iPSC-based systems and rescued oligodendroglial and behavioral abnormalities in AS mouse models. Notably, ESRβ signaling primarily regulates OPC homeostasis rather than directly driving oligodendrocyte differentiation, pinpointing a stage-specific therapeutic entry point. These findings highlight a tractable oligodendroglial pathway downstream of UBE3A with translational potential for AS.

**One Sentence Summary:** UBE3A regulates oligodendroglial homeostasis for myelination via estrogen receptor-β signaling as a therapeutic target for Angelman syndrome.

## INTRODUCTION

UBE3A, also known as E6-associated protein (E6AP) (*1*), is a multifunctional E3 ubiquitin ligase essential for brain development and involved in various signaling pathways beyond its ubiquitin ligase activity (*2–4*). UBE3A is located within the imprinted 15q11–13 locus and displays striking cell-type–specific regulation in the brain: in most mature neurons, UBE3A is predominantly expressed from the maternal allele with the paternal allele silenced (*5*), whereas in glial lineages, including oligodendroglial cells, UBE3A is expressed from both alleles (*6–8*). Consistent with the critical role of UBE3A in neurodevelopment, maternal loss-of-function mutations in UBE3A cause Angelman syndrome (AS), a severe neurodevelopmental disorder characterized by intellectual disability, epilepsy, and a broad spectrum of behavioral phenotypes (*9, 10*). In experimental settings where both UBE3A alleles are disrupted, phenotypes are more severe (*7, 11*), consistent with a greater overall loss of UBE3A dosage that extends beyond classic neuron-restricted imprinting effects. Together, these imprinting and cell-type-specific expression features underscore the complexity of UBE3A biology in the developing brain and motivate investigation of how reduced UBE3A impacts not only neurons but also glial lineages that support circuit maturation. Overall, these observations highlight the importance of maintaining UBE3A within an appropriate dosage range across neural cell types during brain development.

Therapeutic strategies for AS have largely focused on restoring neuronal UBE3A by unsilencing the paternal allele. While preclinical studies have provided important proof-of-concept (*12, 13*), the translation of these approaches to sustained clinical benefit remains an active area of ongoing investigation. This may reflect both the complexity of imprinting biology and the possibility that non-neuronal cell types contribute to disease-relevant circuitry and behavior. Consistent with this view, multiple clinical programs targeting paternal-allele unsilencing have advanced into human testing, with variable trajectories across candidates to date (*14–16*). In parallel, accumulating evidence points to abnormalities in white matter development and myelination in AS, motivating investigation into oligodendroglial mechanisms that could complement neuron-directed approaches (*17–23*). Critically, however, delineating the relative contributions of neuronal versus oligodendroglial UBE3A loss - and defining how neuron–glia interactions shape these phenotypes - remains an open problem that must be addressed to guide therapeutic development.

Oligodendrocyte precursor cells (OPCs) are among the most prolific proliferators in the central nervous system (CNS) (*24*). Their relatively uniform density across many postnatal brain regions suggests robust homeostatic regulation that supports timely production of myelinating oligodendrocytes (*25–27*). Most established models emphasize extrinsic control of OPC proliferation by neighboring cells and secreted cues (*28, 29*), with PDGF-AA/PDGFRα signaling recognized as a major axis supporting OPC proliferation and population maintenance (*30–32*).

However, emerging *in vivo* genetic and quantitative analyses indicate that OPC dynamics also reflect intrinsic, constitutive programs. Recent work shows that OPCs continue to initiate differentiation throughout the adult CNS with stereotyped kinetics that are largely uncoupled from myelin demand or oligodendrocyte loss, instead declining with aging and acute inflammation (*33*). These observations, together with evidence that disease states can disrupt OPC homeostasis - such as reduced OPC density or impaired regenerative responses in chronic demyelination (*34, 35*), and focal OPC hyperdensity linked to functional impairment in select neurodevelopmental disorders (*36*) - point to additional mechanisms, including cell-intrinsic regulators, that shape OPC homeostasis and ultimately influence myelin-dependent circuit function.

Here, we tested whether UBE3A governs OPC self-renewal through an intrinsic mechanism that supports oligodendroglial homeostasis. Using human iPSC-derived OPCs (iOPCs), we found that UBE3A deficiency reduces OPC proliferation and leads to downstream myelination defects, independent of PDGFRα signaling. To identify tractable pathways downstream of UBE3A, we performed a targeted compound screen in UBE3A-deficient iOPCs and discovered that selective estrogen receptor-β (ESRβ) agonists compensate for impaired OPC self-renewal, implicating disrupted ESRβ signaling. We validated these findings in AS patient–derived iPSCs and in AS mouse models, observing reduced ESRβ expression/signaling in AS OPCs and demonstrating that ESRβ agonism restores oligodendroglial homeostasis and improves learning-related behaviors. In parallel, and to directly address cell-type causality and neuron–oligodendroglial interactions, we incorporated oligodendrocyte-lineage–restricted in vivo Ube3a depletion and reductionist neuron–OPC co-culture paradigms, supporting an intrinsic requirement for oligodendroglial UBE3A in OPC proliferation and myelination while clarifying the contribution of neuronal context. Collectively, these results establish a UBE3A-dependent pathway that regulates OPC homeostasis and nominate ESRβ signaling as a stage-specific, therapeutically actionable entry point that can complement ongoing neuron-directed UBE3A restoration strategies in AS.

## RESULTS

### UBE3A loss in AS reduces oligodendrocyte populations and myelination by impairing OPC proliferation

We first confirmed whether the depletion of UBE3A corresponds to the oligodendroglial dysfunction observed in AS mice *in vivo*. We choose to use the established and well-characterized AS mouse model (*Ube3a^m−/p+^,* or E6-AP *Ube3a^tm1Alb^*) (*5*), which well recapitulates key features of neuronal imprinting and behavioral abnormalities. *Ube3a* imprinting is cell type-specific across the CNS: in mature neurons broadly. UBE3A is predominantly expressed from the maternal allele with the paternal allele silenced, such that maternal loss produces widespread neuronal UBE3A deficiency throughout the brain (*5*). In contrast, in non-neuronal tissues *Ube3a* is generally expressed more biallelically, and accordingly maternal *Ube3a* deficiency mice exhibit haploinsufficiency in peripheral organs, including a ∼70–80% reduction in UBE3A protein levels in heart, liver, and kidney (*37*). The even more severe reduction in UBE3A expression in almost all regions of the brain is associated with microcephaly in AS, a highly penetrant syndromic feature with early postnatal onset (*38*). To explore oligodendroglial population dynamics and myelination across development, we examined multiple oligodendroglial markers in brain sections from AS mice at early postnatal timepoints and into maturity. Because myelination changes rapidly during the first postnatal weeks, we analyzed the early postnatal stage as separate P7 and P10 cohorts, and compared these with juveniles (P30) and adults (P180). While differences across these age groups were evident, the overall deficit in AS relative to WT was clear at each stage. In agreement with reports in the literature (*39*), we first noticed in AS pups a downsized pool of oligodendroglia, accompanied by decreased myelin expression and a shrinkage of gray and white matter structures, in the brain regions of the hippocampus (**Fig. 1A-B**) and the corpus callosum and fimbria of the fornix (**Fig. S1A-B**). In juvenile and adult AS mouse brains, we observed a significant decline in oligodendrocyte populations (**Fig. 1B**) and a moderate yet distinct decrease in myelin expression (**Fig. S1C**). This was associated with consistent thinning of the cortex and corpus callosum up to six months of age (**Fig. S1D**). Of note, we further supported these findings by reanalyzing an independent, published cortical proteomics dataset from a distinct AS transgenic model and observing reduced abundance of multiple oligodendrocyte- and myelination-associated proteins (*40*) (**Fig. S2**). Furthermore, UBE3A expression was significantly reduced in oligodendroglia across various developmental stages in AS mice compared to wild-type controls (**Fig. 1B**). These findings support our hypothesis that reduced UBE3A expression in the oligodendrocyte lineage disrupts oligodendroglial homeostasis, contributing to myelination deficits during neurodevelopment.

**Figure 1.**
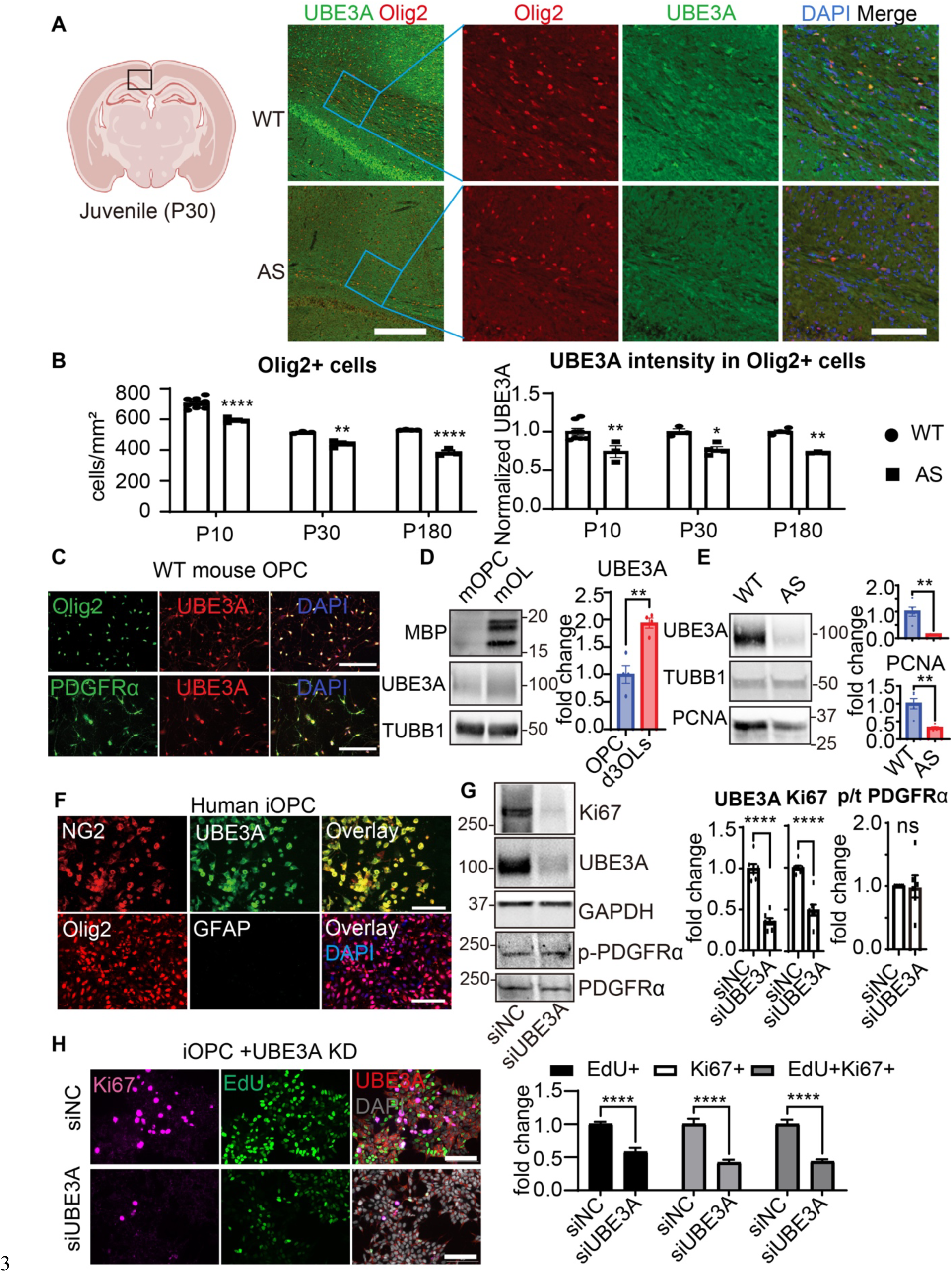
UBE3A expression across the oligodendrocyte lineage supports OPC proliferation in AS-relevant mouse and human models. (A) Assessment of oligodendroglial populations and UBE3A protein levels in hippocampal CA1 sections of WT and AS juveniles at postnatal day 30 (P30). Oligodendrocytes were identified with Olig2 (red), UBE3A was labeled in green, and nuclei were counterstained with DAPI (blue). Scale bars: left, 100 µm; right, 50 µm. (B) The oligodendroglial cell density and UBE3A expression were quantified across developmental milestones: pups (P10), juveniles (P30), and adults (P180). The data reflect the number of Olig2-positive oligodendrocytes per area, with UBE3A intensity normalized against wildtype levels (set at 1.0). n=3-8 animals per group. (C) Immunofluorescence validation of UBE3A expression in purified mouse primary OPCs, identified by PDGFRα and Olig2, with DAPI nuclear counterstain. Scale bar 200 um. (D) Immunoblot showing MBP upregulation during differentiation of WT mouse primary OPCs into oligodendrocytes. UBE3A and MBP were quantified and normalized to TUBB1. n = 4 pups from 2 independent experiments. (E) Immunoblot analysis of primary OPC proliferation from WT and AS mice, assessed by PCNA. UBE3A and PCNA were quantified and normalized to TUBB1. n = 6-10 pups from 3 independent experiments. (F) Immunofluorescent staining to validate UBE3A expression in iPSC-derived oligodendrocyte precursor cells (iOPCs) marked by NG2, Olig2 and DAPI, but devoid of negative control GFAP. Scale bar 100 um. (G) Immunofluorescent staining to assay the OPC proliferation altered by UBE3A knockdown, measuring the expression of two independent proliferation markers, Ki67 (magenta) expression and EdU (green) nuclear incorporation, in iOPCs treated with non-targeting control siRNA (siNC) or siRNA against UBE3A (siUBE3A) and labeled with DAPI (blue) and UBE3A (red); data were plotted by cell numbers positive for either or both markers (normalized to control, siNC=1.0). n=40-48 regions of interest from 3 independent experiments. (H) Immunoblotting for measurement of Ki67 expression and PDGFRα activation (Y754 phosphorylation) in iOPCs subject to RNAi-mediated UBE3A knockdown by siNC or siUBE3A. n=7 experiments. Scale bar 100 um.

To test whether reduced UBE3A is linked to oligodendroglial dysfunction through a cell-intrinsic mechanism, we acutely isolated a highly enriched population of oligodendrocyte precursor cells (OPCs) from the developing mouse brain and verified their appropriate lineage identity (**Fig. 1C**). When these primary OPCs were induced to mature *in vitro*, these oligodendrocytes robustly engaged a myelin-associated differentiation program (**Fig. 1D**), validating their functional competence. Importantly, OPCs isolated from the AS mouse brains showed lower UBE3A levels together with reduced proliferative capacity, supporting the idea that UBE3A is required to sustain the proliferative pool that ultimately supplies cells for myelination (**Fig. 1E**).

We next asked whether UBE3A directly governs the self-renewal behavior of human OPCs in a reductionist iPSC-derived system, a model that has been well documented to be appropriate for studying how oligodendrocytes mature and form myelin(*41–43*). We developed and optimized a chemically-defined protocol(*44, 45*), which demonstrated a full yet accelerated development course for monitoring a wide spectrum of oligodendroglial functions (**Fig. S3A**) – from the stages of neural precursor cells (NPCs; Week 1), oligodendrocyte precursor cells (OPCs, Week 2), pre-myelinating oligodendrocytes (pre-OLs, Week 3) to mature oligodendrocytes (OLs, Week 4-5). The resultant iOPCs possessed proper morphology, expressed specific functional markers (**Fig. 1F, S3A**), and could be further differentiated into mature iOLs for the in vitro myelination assay. Notably, UBE3A expression was detected in all oligodendroglial development stages; and its level was most notable in OPCs and pre-OLs (**Fig. S3B**), consistent with the independent results of single-cell oligodendroglial transcriptomics on the developing mouse brains(*46*) (**Fig. S3C**). The pattern of UBE3A expression we identified points to its significant role during the early stages of oligodendrocyte differentiation. Given the inherent dynamic quality of oligodendrocyte progenitor cells (OPCs), which continually self-renew to preserve their population via local homeostatic proliferation(*47*) – making them highly sensitive to environmental and genetic changes, we aimed to examine this process more closely.

To elucidate the specific role of UBE3A in OPC self-renewal and subsequent differentiation into myelinating oligodendrocytes, we employed targeted siRNAs to reduce UBE3A expression in iPSC-derived OPCs (iOPCs). The effectiveness of this reduction was demonstrated by a series of siRNAs achieving over 50% knockdown efficiency (**Fig. 1G**). This significant reduction in UBE3A levels corresponded with a notable decrease in iOPC proliferation, as indicated by lower expressions of the proliferation marker Ki67 and reduced EdU incorporation into DNA (**Fig. 1G-H**), though it did not trigger cell death (**Fig. S3D**). To validate these findings, primary OPCs were isolated from rat brains(*48, 49*) and subjected to similar UBE3A knockdown, which similarly curtailed OPC proliferation without causing cell death (Fig. **S3E-F**). Notably, this impaired proliferation was not linked to changes in PDGFRα receptor activation (**Fig. 1G**). Moreover, a direct correlation was established between diminished UBE3A protein levels and reduced iOPC proliferation, where even a moderate reduction of UBE3A (20-40%) significantly impeded cell proliferation (**Fig. S3G**). This supports our hypothesis that UBE3A functions as an intrinsic cellular regulator in OPCs, and OPC proliferation is acutely sensitive to variations in UBE3A gene dosage, with even moderate decreases such as those observed in AS with biallelic expression impacting OPC homeostasis.

### Cell-intrinsic requirement for UBE3A in the oligodendroglia revealed by lineage-restricted in vivo knockdown

Because AS mice carry neuronal UBE3A loss that can indirectly influence oligodendroglial development and myelination through altered circuit activity and trophic support, *in vivo* phenotypes observed in the maternal-deficiency model could reflect a mixture of cell-intrinsic oligodendroglial requirements and non-cell-autonomous neuronal effects (*51–54*). To disentangle these contributions and directly test the necessity of UBE3A within the oligodendrocyte lineage in an otherwise wild-type brain, we developed an oligodendrocyte-lineage–restricted knockdown strategy using neonatal AAV delivery. We chose an Olig1 promoter–driven rAAV-Olig001 platform because it has been shown to provide strong, preferential CNS oligodendrocyte targeting with minimal off-target/peripheral transduction in rodents (*55, 56*), and related oligodendrocyte-targeting AAV vectors have advanced to intracranial gene therapy clinical trials (*57*). Specifically, wildtype neonatal pups received intracerebroventricular injection at P0 of an AAV expressing either a non-targeting control shRNA or one of two independent shRNAs targeting *Ube3a* under an oligodendroglial promoter, with GFP marking transduced cells (**Fig. 2A**). We then used a staged experimental design: a subset of mice was harvested at P10 to validate lineage-restricted targeting and knockdown efficiency and to biochemically assess proliferative state in purified OPCs, while the remaining mice were tested for motor learning at P40 and harvested at P42 for histological analyses of proliferation and myelination (**Fig. 2A**).

**Figure 2.**
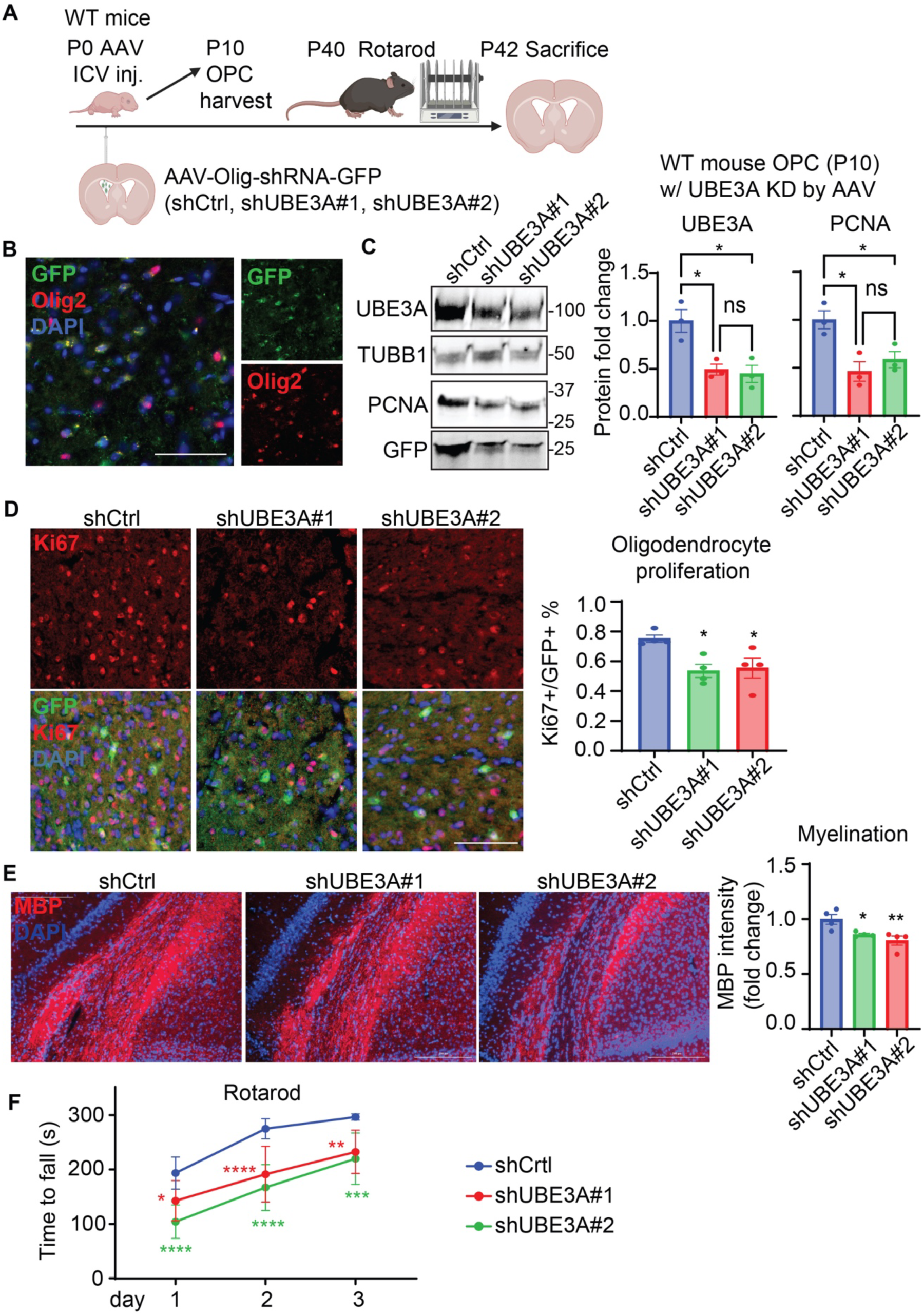
Oligodendrocyte-lineage–restricted depletion of UBE3A impairs OPC proliferation, myelination, and motor learning *in vivo*. **(A)** Experimental schematic for oligodendrocyte-lineage–restricted UBE3A knockdown *in vivo*. WT neonatal pups received P0 intracerebroventricular injection of AAV-Olig1-shRNA-GFP expressing a non-targeting control shRNA (shCtrl) or two independent shRNAs targeting UBE3A (shUBE3A#1 and shUBE3A#2; distinct target regions), with GFP marking transduced cells. A subset of animals was harvested at P10 for validation of transduction specificity and knockdown efficiency and for primary OPC purification. Remaining animals underwent rotarod testing at P40 and were harvested at P42 for histological assessment of proliferation and myelination. **(B)** Validation of oligodendrocyte-lineage transduction by AAV-Olig1–shRNA-GFP. Viral expression was detected by GFP (green), oligodendrocyte-lineage cells were labeled by Olig2 (red), and nuclei were counterstained with DAPI (blue). Scale bar, 200 µm. **(C)** Immunoblot analysis of primary OPCs purified from P10 AAV-injected mice showing UBE3A knockdown and altered proliferation marker expression. UBE3A and PCNA were quantified and normalized to TUBB1. *n* = 4 pups. **(D)** *In vivo* OPC proliferation at P42 assessed by immunofluorescence for Ki67 (red) within GFP+ transduced cells (green), with DAPI nuclear counterstain (blue). Quantification shown as the percentage of Ki67+ cells among GFP+ cells. *n* = 4 animals per group. **(E)** Myelination at P42 assessed by immunofluorescence for MBP (red) with DAPI nuclear counterstain (blue). Quantification shown as MBP intensity normalized to the GFP control condition. *n* = 4 animals per group. **(F)** Motor coordination and learning assessed by the rotarod test (P40–P42), plotted as latency to fall across training days for shCtrl and shUBE3A groups. *n* = 9–10 mice per group.

We first validated the knockdown reagents and early *in vivo* targeting. Both *Ube3a*-directed shRNAs robustly reduced UBE3A protein in a cell-based assay (**Fig. S4A**), supporting their use for *in vivo* studies. Following P0 injection, we confirmed that GFP expression was preferentially localized to oligodendrocyte-lineage cells in the developing brain (**Fig. 2B**). At P10, we purified OPCs from injected mice and found that UBE3A was reduced in the UBE3A shRNA groups relative to control, demonstrating efficient oligodendroglial knockdown *in vivo* (**Fig. 2C**). Importantly, this reduction was accompanied by a decrease in a proliferation-associated marker in purified OPCs, providing early biochemical evidence that oligodendroglial UBE3A supports the proliferative state of the OPC pool (**Fig. 2C**).

We next assessed the longer-term consequences of oligodendrocyte-lineage UBE3A depletion *in vivo*. Imaging of the injected hemisphere showed widespread GFP signal across major forebrain regions, with preferential labeling of oligodendrocyte-lineage cells in areas such as corpus callosum, external capsule, and striatum (**Fig. S4B-E**). We further verified knockdown at the cellular level by quantifying UBE3A immunofluorescence intensity specifically within shRNA-transduced GFP+ cells, which confirmed a significant reduction in UBE3A signal *in vivo* (**Fig. S4F**). Critically, GFP expression showed minimal overlap with markers of astrocytes, neurons, or microglia, supporting that the knockdown was largely restricted to the intended lineage (**Fig. S4G**). At P42, this oligodendrocyte-lineage UBE3A depletion led to a measurable reduction in proliferation among shRNA-expressing cells *in vivo* (**Fig. 2D**), and was sufficient to reduce myelin marker intensity in brain tissue (**Fig. 2E**), closely phenocopying the myelination deficits observed in AS mice and reinforcing a cell-intrinsic requirement for UBE3A in sustaining oligodendroglial output.

Because myelination is increasingly recognized as a determinant of circuit performance, prior studies have shown that oligodendrocyte-lineage-specific genetic perturbations that disrupt the supply of new myelinating cells can produce measurable deficits in motor learning and cognition, supporting a causal link between oligodendroglial dysfunction and behavior (*53, 54, 58*). We therefore asked whether oligodendrocyte-lineage UBE3A depletion alone is sufficient to elicit a behavioral phenotype relevant to AS. Among commonly used behavioral paradigms, rotarod performance is particularly sensitive to myelin and oligodendrocyte disruption, including in multiple models of demyelinating disease (*59, 60*). Moreover, across multiple independent studies using standardized protocols, the accelerating rotarod assay yields a robust and reproducible deficit in maternal UBE3A-deficient mice and is widely regarded as one of the most reliable behavioral readouts in AS models (*61–64*). Consistent with this rationale, mice receiving oligodendrocyte-lineage UBE3A knockdown exhibited significantly impaired rotarod learning compared to controls (**Fig. 2F**). This phenotype mirrors a key behavioral deficit in AS models and aligns with evidence that compromised oligodendroglial development and myelin integrity can limit experience-dependent learning. Together, these findings provide *in vivo* evidence - independent of neuronal UBE3A loss - that UBE3A within the oligodendrocyte lineage is required to sustain OPC proliferation, support myelin output, and enable normal motor learning.

### The activation of estrogen receptor-β (ESRβ) function mitigates the OPC proliferation impaired by UBE3A depletion

Given the scarcity of knowledge regarding the role of UBE3A in the biology of oligodendrocytes, we systemically explored a broad array of cellular functions, utilizing the high-throughput capabilities of our iPSC-based methodology. Recognizing the pronounced change in OPC proliferation in response to UBE3A expression levels, we carefully chose over 45 compounds known for their targeted cellular actions (**Table S1**). These compounds were meticulously selected based on literature that either connected them directly to UBE3A, noted their ability to promote oligodendrocyte growth or maturation, or documented their protective effects on brain tissue. We then fine-tuned the reduction of UBE3A using siRNA to achieve a roughly 50% decrease in OPC proliferation, monitored by Ki67 protein levels (**Fig. 3A**). This reduction set the stage for us to test the individual compounds, gauging their potential to reverse the proliferation impairment caused by UBE3A depletion and to uncover underlying cellular mechanisms responsible for the observed deficits.

**Figure 3.**
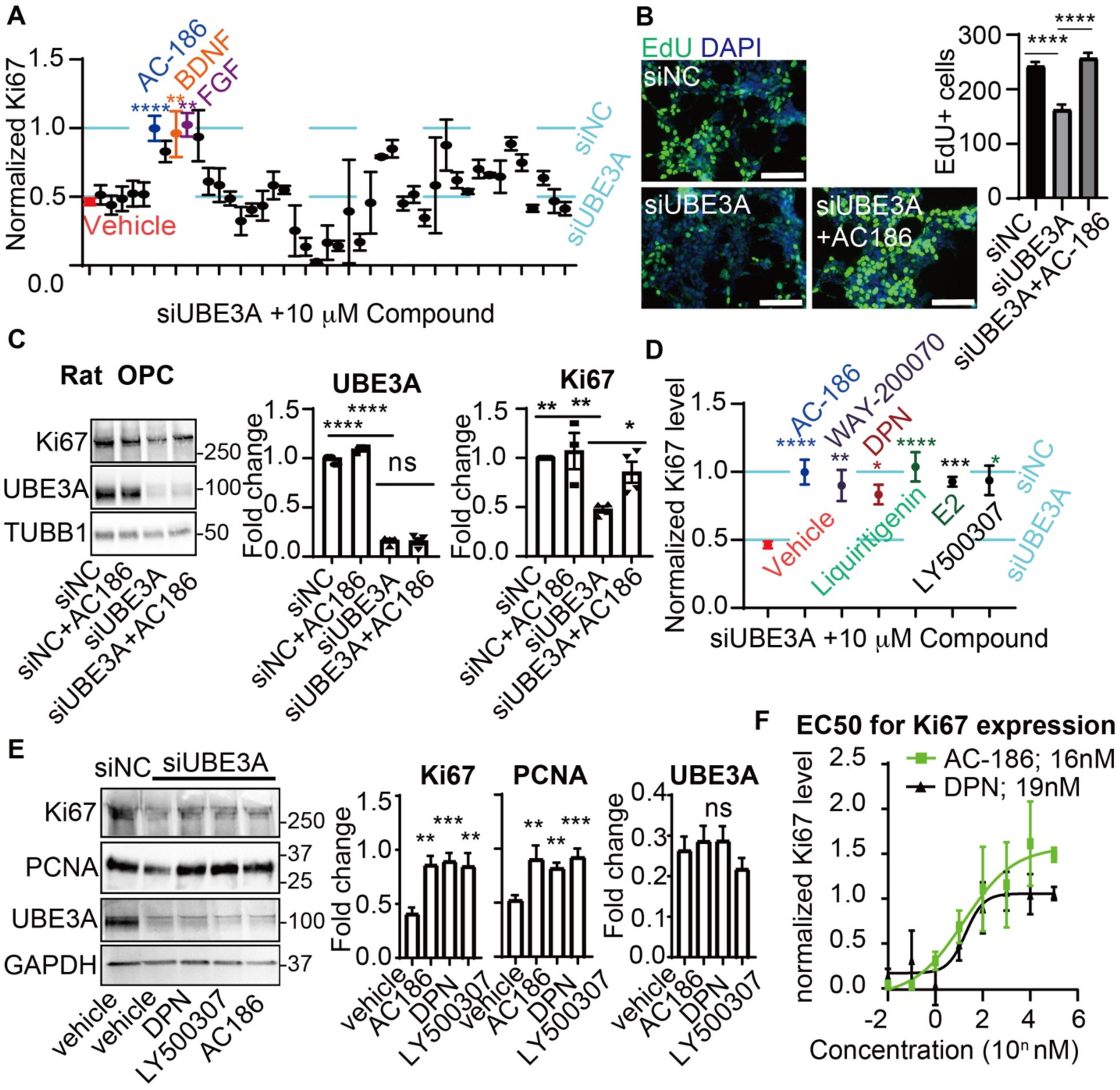
Identification of estrogen receptor-β (ESRβ) agonists to mitigate OPC proliferation deficits from UBE3A depletion. **(A)** Compound screening identified agents capable of restoring OPC proliferation reduced by UBE3A loss. Forty-five compounds (listed in **table S1**) were individually tested at 10 µM over 24 hours on UBE3A-deficient iOPCs with around a 50% reduction in Ki67 expression. The potential of these compounds to reinstate Ki67 expression, relative to vehicle control (DMSO), was quantified via immunoblotting, with Ki67 levels normalized to beta-tubulin and compared to the siNC group (value=1.0). AC-186, a highly selective estrogen receptor-β (ERβ) agonist (blue); BDNF, brain-derived neurotrophic factor (orange); FGF, fibroblast growth factor (purple). n=3-10 per condition. **(B)** Immunofluorescence to confirm the impact of AC-186 on OPC proliferation, with EdU incorporation in iOPC nuclei measured across three conditions: siNC plus vehicle, siUBE3A plus vehicle, and siUBE3A plus AC-186 at 10 µM for 24 hours. Cell counts positive for EdU were normalized to the siNC group (siNC=1.0), with n=24-28 region of interest from 3 independent experiments. **(C)** Cross-platform validation of the efficacy of AC-186 using rat primary OPCs deficient in UBE3A treated with siNC or siUBE3A and either vehicle or AC-186, 10 µM over 24 hours, across four conditions. Densitometric analysis of UBE3A and Ki67, normalized to β-tubulin (TUBB1), was compared to siNC plus vehicle (value=1.0), with n=4 per condition. **(D)** Assessment of multiple ESRβ agonists with distinct chemical structures, including AC-186, WAY-200070, DPN, LY500307, Liquiritigenin and E2 (**Table S1**), for their restorative effects on OPC proliferation. Each compound’s selectivity was noted, especially the strong preference of AC-186 for ESRβ over ESRα, with n=3-9 per condition. **(E)** Efficacy of ESRβ agonists in mitigating proliferation deficits in UBE3A-depleted OPCs was verified by immunoblotting, assessing the levels of 2 distinct proliferation markers, Ki67 and PCNA, post-treatment with AC-186, DPN, and LY500307 (10 µM, 24 hours). Protein levels were normalized to GAPDH and compared to siNC plus vehicle (set at 1.0). n=4-9 per group. **(F)** Establishment of the half-maximal effective concentration (EC50) for AC-186 and DPN, which reflects the potency of these compounds in reversing the proliferation deficits of UBE3A-deficient OPCs, by applying a range of drug concentrations to iOPCs treated with siUBE3A for 24 hours. Ki67 expression was measured as a marker of proliferation as the immunoblotting. The EC50 calculations, which indicate the concentration at which each drug achieves a 50% maximal response, were derived from the dose-response curves. These curves plot Ki67 expression levels, normalized to the siNC plus vehicle group (set as 1.0), against drug concentrations. The EC50 values were determined to be approximately 16 nM for AC-186 and 19 nM for DPN, revealing their relative effectiveness in a cellular context mimicking UBE3A loss.

Our screening identified specific compounds, including brain-derived neurotrophic factor (BDNF) (*65, 66*), fibroblast growth factor (FGF)(*67, 68*) and AC-186 (*69–71*), that effectively normalized proliferation levels in UBE3A-deficient iOPCs (**Figure 3A-B**). AC-186 was particularly notable for its high specificity to ERβ and its proven efficacy in neurological disorders (*69, 71*). To rigorously validate our results across different platforms, we isolated primary OPCs from neonatal rat brains (*48, 49*) and replicated the reduction in proliferation by RNAi-mediated UBE3A depletion. Similarly, the proliferation inhibition was reversed by AC-186, as observed in human iOPCs (**Figure 3C**). To ensure these effects were specific and not due to off-target actions, we compared AC-186 with other selective ERβ activators of different structural classes(*72*) (**Table S1**), including WAY-200070, diarylpropionitrile (DPN), WAY-200070, LY500307, Liquiritigenin, and estradiol (E2); all of which similarly mitigated the impact of UBE3A loss on OPC proliferation (**Figure 3D-E, S1G**). Additionally, to address potential artifacts from siRNA-based knockdown, we utilized CRISPR/Cas9 technology to decrease UBE3A expression with multiple sgRNAs, confirming the consistent impact of UBE3A depletion on OPC proliferation and the therapeutic effects of ERβ agonists on impaired OPC proliferation (**Fig. S5A**). Notably, neither AC-186 nor DPN affected human or rat OPCs with normal UBE3A levels (**Fig. S5A-B**), highlighting their specific action in conditions associated with UBE3A deficiency. We determined effective concentrations (EC50) for AC-186 and DPN to restore Ki67 expression to be approximately 16 nM and 19 nM, respectively (**Fig. 3F**). Collectively, these results identify ERβ activation as a selective and reproducible strategy to restore the self-renewal deficit caused by UBE3A depletion, motivating us to next determine which cell type(s) mediate the functional rescue in the context of neuron-oligodendrocyte interactions.

To directly dissect cell-intrinsic versus interaction-dependent contributions, we turned to a reductionist human iPSC-based neuron-oligodendroglial co-culture model (*44, 45*) in which control or UBE3A-deficient induced neurons were paired with control or UBE3A-deficient iOPCs and differentiated toward a myelinating state. Across these four combinations, we quantified myelin output as a readout of oligodendroglial function and found that neuronal UBE3A deficiency measurably reduced myelination, consistent with a non-cell-autonomous influence of neurons on the myelinating program. Importantly, however, oligodendroglial UBE3A deficiency exerted a clear intrinsic effect: co-cultures containing UBE3A-deficient oligodendroglial cells displayed similarly reduced myelination regardless of whether neurons were control or UBE3A-deficient, whereas control neuron + control oligodendroglial co-cultures showed the highest myelin output (**Fig. S6A**). Notably, treatment with AC-186 elevated myelin output across all four conditions to a common level comparable to vehicle-treated control neuron + control oligodendroglial cultures (**Fig. S6A**), supporting the conclusion that ERβ agonism primarily restores oligodendroglial function rather than acting through neurons to enhance myelination.

In parallel, we asked whether AC-186 could modulate a well-established neuronal functional deficit in AS, namely impaired synaptic plasticity. Long-term potentiation (LTP) is consistently disrupted in AS models and serves as a canonical readout of synaptic dysfunction (*5, 73*); importantly, accumulating evidence also links effective LTP to intact oligodendrocyte support and myelination (*74, 75*). We therefore assayed synaptic plasticity using a chemical LTP paradigm (glycine stimulation) and quantified phosphorylation of an AMPA receptor subunit as a molecular proxy for LTP-related signaling (*76, 77*). In induced neurons cultured alone, UBE3A depletion reduced the plasticity-associated phosphorylation signal, and AC-186 did not rescue this deficit (**Fig. S6B**), arguing against a direct neuronal mechanism of action. Strikingly, when neurons were co-cultured with oligodendroglial cells under myelinating conditions, AC-186 restored the plasticity-associated phosphorylation deficit in the setting of UBE3A depletion (**Fig. S6B**). Together, these findings indicate that ERβ agonism improves neuronal plasticity readouts indirectly, through restoration of oligodendroglial function and myelination, and further support a model in which UBE3A loss in oligodendroglial cells drives a primary deficit in the self-renewing/myelinating program that can secondarily impact neuronal function.

### UBE3A depletion suppresses ESR-β signaling through MAPK/ERK pathway in OPC self-renewal

The effectiveness of AC-186 in reversing the impaired OPC proliferation due to UBE3A depletion directed our focus to the role of estrogen receptor signaling in oligodendroglia. Estrogen receptors are prevalent in the brain, found in both neuronal and glial cells and are essential for a variety of cellular functions independent of gender (*78*). Our study confirmed that all three known receptors are expressed in iOPCs (**Fig. 4A-B**): estrogen receptor-α (ESR-α), -β (ESR-β) and G protein-coupled estrogen receptor (GPER). Activation of estrogen receptors, either by ligands (e.g. estrogen or AC-186) or by a ligand-independent mechanism involving cofactors (*79–81*), can engage canonical signaling cascades that regulate proliferation and differentiation. Notably, UBE3A knockdown in iOPCs produced a coordinated shift in receptor profiles: GPER was upregulated while ESR-β was reduced (**Fig. 4B**), consistent with a selective vulnerability of the ESR-β axis in the oligodendroglial lineage and aligning with the pharmacologic specificity of AC-186. In contrast, in iPSC-derived human neurons (*82, 83*), UBE3A knockdown resulted only in increased GPER levels without affecting ESR-α or ESR-β levels (**Fig. S7A-B**). UBE3A depletion likewise did not change estrogen receptor expression in iPSC-derived astrocytes (*84*) or microglia (*85*) (**Fig. S7A-B**). Importantly, our compound screen (**Fig. 3A**) included a selective GPER ligand (G-1; **Table S1**), which did not significantly rescue the OPC proliferation deficit under our screening conditions, supporting our prioritization of ESR-β agonism for mechanistic and translational follow-up.

**Figure 4.**
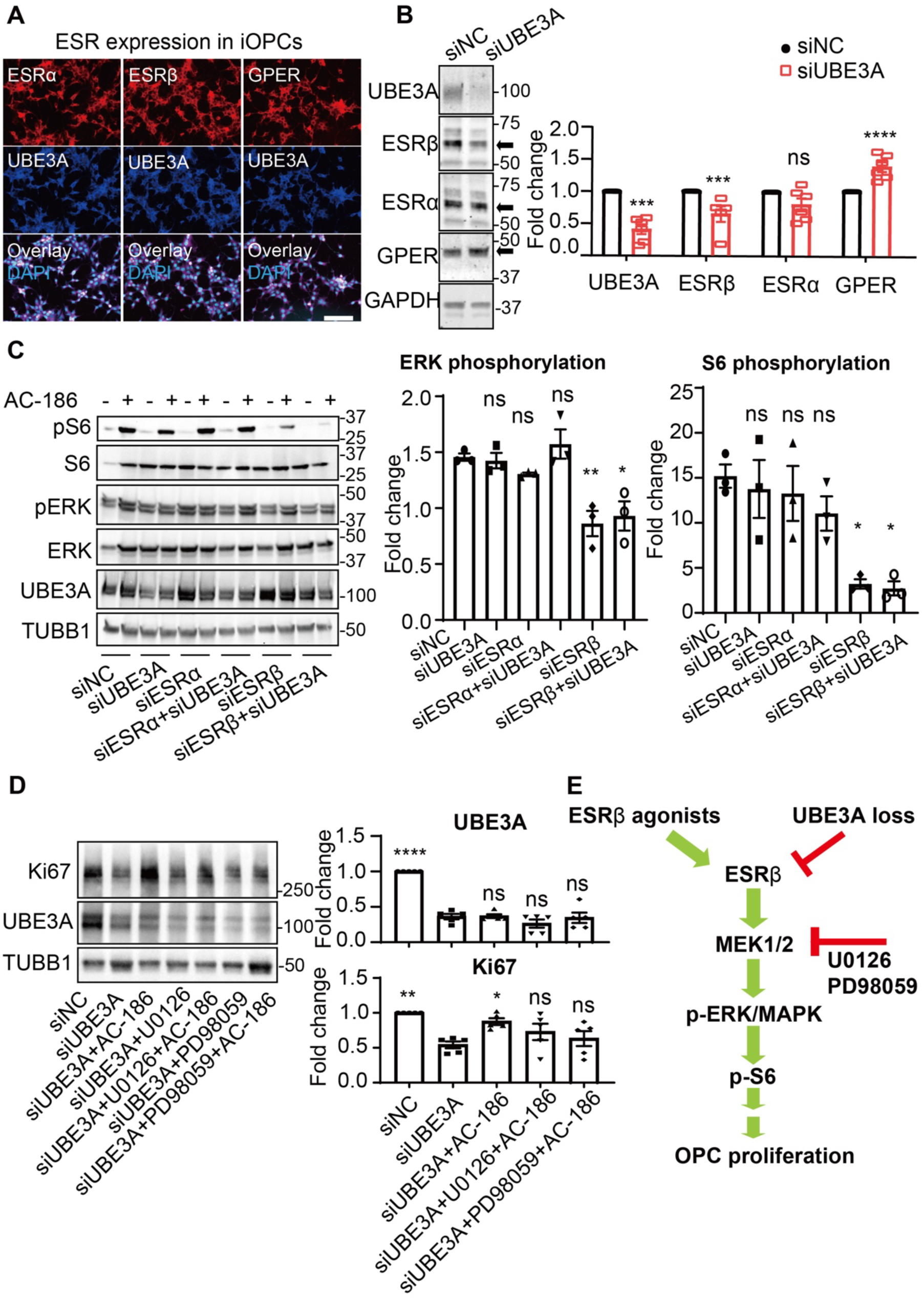
Identification of ESR-β signaling pathway in regulating OPC proliferation impaired by UBE3A depletion. **(A)** Immunofluorescence to track estrogen receptor (ESR) expression in OPCs, by staining of all 3 ESRs, ESR-α, ESR-β and GPER (red), in iOPCs co-labeled with UBE3A (blue) and DAPI (light blue); scale bare, 100 μm. **(B)** Immunoblotting to assess OPC expression of individual ESRs, by analyzing protein levels of ESR-α, ESR-β and GPER in iOPCs treated with siNC or siUBE3A, normalized to GAPDH and expressed relative to the siNC condition (set as 1.0). n=5-6 experiments. **(C)** The activation of the ERK/MAPK signaling pathway in OPCs, particularly downstream of ESR-β, was analyzed in iOPCs subjected to 6 knockdown conditions: siNC, siUBE3A, siESRα, siESRβ, and combinations of siUBE3A with siESRα or siESRβ. These were followed by treatment with vehicle control or the ESR-β agonist AC-186 (10 µM, 30 min). The activation of ERK and its substrate S6 was determined by the ratio of phosphorylated to total ERK and S6 levels, adjusted relative to the vehicle control group (standard value of 1.0). n=3 experiments. **(D)** Immunoblotting to evaluate the requirement for ERK/MAPK activation in the restoration of OPC proliferation following ESR-β activation, with the addition of ERK phosphorylation inhibitors U0126 or PD98059 to UBE3A-depleted OPCs, with or without AC-186 co-treatment (10 µM, 24 hours). The quantification of Ki67 expression and UBE3A levels, being normalized to β-tubulin (TUBB1) and compared to the siNC condition (value of 1.0). n=5 experiments. **(E)** A schematic depicting the regulatory role of ESR-β signaling pathway in OPC proliferation, identified by combining genetic knockdown techniques and pharmacological interventions. Activation of ESR-β selectively triggers the ERK/MAPK pathway, as shown by ERK phosphorylation and the ERK substrate S6 activation, essential for cell proliferation. Reduction in ESR-β function due to UBE3A loss can be compensated by ESR-β agonists, suggesting a mechanism to improve impaired OPC proliferation. Statistical significance was determined using one-way ANOVA followed by post hoc multiple-comparison testing, with pre-specified pairwise comparisons displayed. For (**B, C**), comparisons were presented against the siNC control condition. For (**D**), selected comparisons were made between siUBE3A and the indicated treatment conditions (e.g., siUBE3A+AC-186, siUBE3A+U0126+AC-186, siUBE3A+PD98059+AC-186). Significance is denoted as * p<0.05, ** p<0.005, *** p<0.001, **** p<0.0001; ns, not significant.

Further exploring the role of estrogen receptor in cellular function, we considered that besides the classical genomic regulatory roles of cytoplasmic and nuclear estrogen receptors (ESR-α and -β), membrane-bound estrogen receptors (ESR-β and GPER) can also initiate rapid intracellular signaling cascades upon activation (*79, 80, 86*). We examined in iOPCs the effects of UBE3A knockdown on multiple pathways downstream of estrogen receptor signaling, including mTOR/4EBP1, ERK/MAPK, PI3K/Akt, TrkB and NFkB. We found a reduction in the activation of the ERK/MAPK pathway and its phosphorylation target, S6, without significant changes in other pathways associated with ESR-β (**Fig. S7C**). Considering the essential role of the ERK/MAPK signaling pathway in the proliferation and differentiation of oligodendroglia (*87*), we sought to determine if the loss of ESR-β was linked to the observed decrease in ERK/MAPK activity. We treated the UBE3A-depleted iOPCs with the ESR-β agonist AC-186 and found that this treatment restored the activity of ERK and its downstream target, S6, while UBE3A levels remained unchanged (**Fig. 4C**). The beneficial effects of AC-186 on ERK/MAPK activation were specifically tied to ESR-β, as demonstrated by the fact that these effects were negated when ESR-β was knocked down. In contrast, knocking down ESR-α, which is structurally similar to ESR-β, did not reverse the effects of AC-186 (**Fig. 4C**). Moreover, the silencing of ESR-β, but not of ESR-α, replicated the consequences of UBE3A loss by diminishing activation in the ERK/MAPK pathway and S6 phosphorylation (**Fig. 4C**). This reinforces the specific involvement of ESR-β in the signaling dysfunction observed in oligodendroglia due to UBE3A deficiency. Additionally, when we introduced the highly selective MEK inhibitors, PD98059 and U0126, they inhibited the ERK/MAPK pathway activation stimulated by AC-186 and consequently reduced the effect of the compound to enhance the proliferation of iOPCs with UBE3A knockdown (**Fig. 4D**).

These observations collectively showed that ESR-β is a key activator of the ERK/MAPK signaling cascade in OPCs, which plays a crucial role in regulating their proliferation (**Fig. 4E**). The disruption of this ESR-β/ERK pathway due to UBE3A depletion could potentially be corrected by the administration of an ESR-β agonist.

### ESR-β downregulation by UBE3A deficiency affects OPC self-renewal but not myelination in oligodendrocytes

Our findings have jointly underscored a critical role for UBE3A in maintaining oligodendroglial homeostasis by regulating cell proliferation via ESR-β signaling. However, the influence of UBE3A on oligodendrocyte differentiation and maturation remains uncertain. We further assessed the mRNA levels of estrogen receptors in iOPCs and noted no alterations following UBE3A depletion (**Fig. S8A**), indicating that regulation of ESRβ expression by UBE3A may occur post-transcriptionally, potentially through mechanisms that stabilize protein levels, as previously documented (*88*). To delve deeper into the effects of UBE3A depletion on oligodendrocyte development, we differentiated iOPCs into pre-myelinating (pre-iOL) and fully mature oligodendrocytes (iOL) (**Fig. S3A-B, S8B-C**), with persistent UBE3A depletion. The UBE3A depletion resulted in a reduced population of mature oligodendrocytes, evidenced by decreased expression of mature oligodendrocyte markers (**Fig. 5A**) and impaired myelination capacity, particularly in their ability to ensheath synthetic nanofibers with myelin (*89*) (**Fig. 5B**). Intriguingly, the downregulation of ESR-β induced by UBE3A depletion was not observed after the differentiation of iOPCs into iOLs (**Fig. 5C**), highlighting a major shift in cellular state.

**Figure 5.**
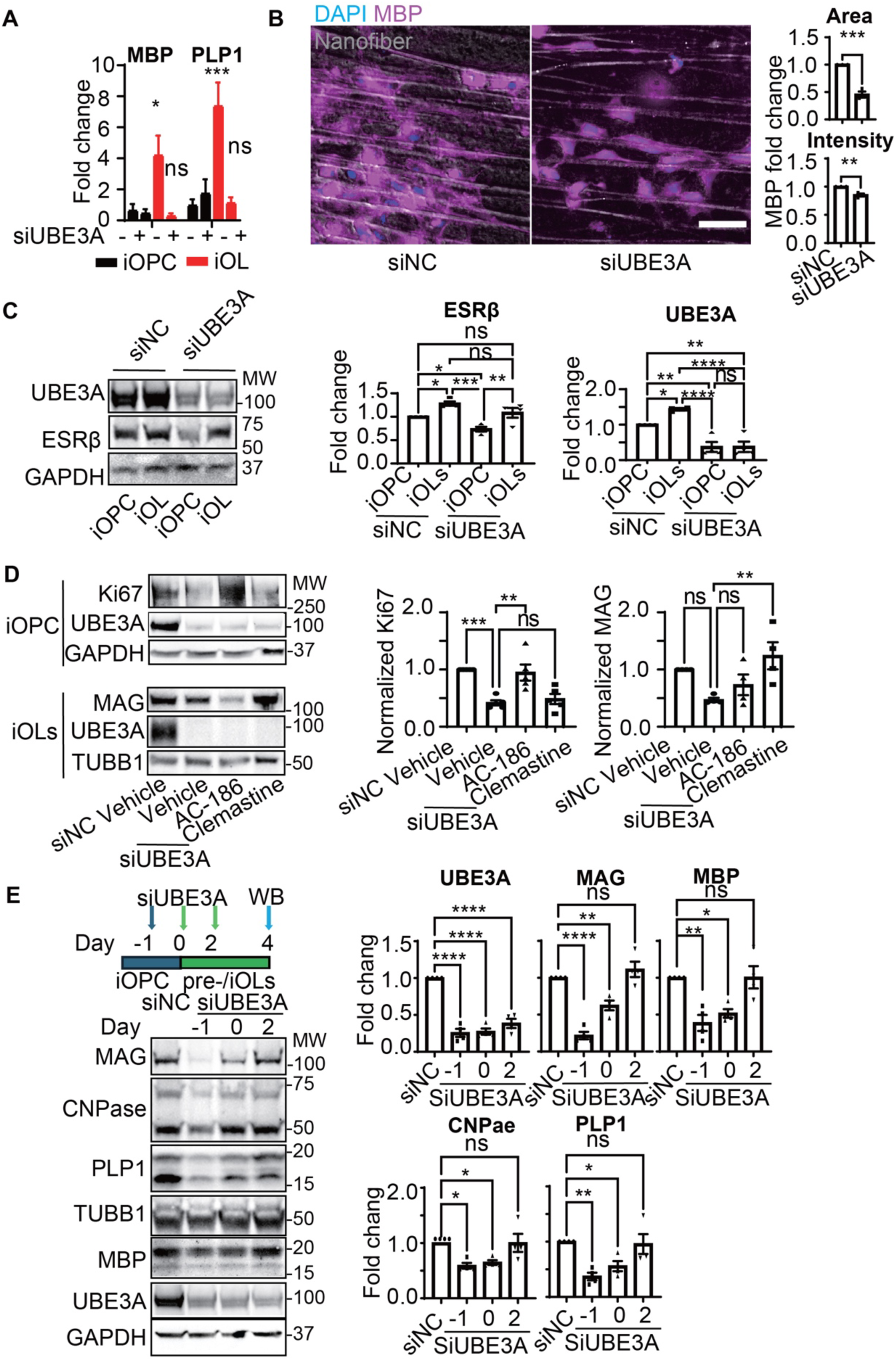
Differential impacts from UBE3A via ESR-β signaling on OPC proliferation and oligodendrocyte differentiation. **(A)** Assessments of qPCR to evaluate the differentiation and maturation for myelination of iOPCs after persistent UBE3A knockdown, by measuring mRNA transcripts of UBE3A and myelination markers MBP and PLP1 in iOPCs treated with siNC or siUBE3A and differentiated *in vitro*. n=3 experiments. **(B)** *In vitro* myelination assays analyze the effect of persistent UBE3A knockdown on iOPCs differentiation and maturation in nano-fiber-based cultures, with immunofluorescent staining of MBP (magenta) and DAPI (blue) and imaging of nano-fibers (grey). Quantification by the area (top graph) and density (bottom graph) of MBP florescent signals (normalized to siNC=1.0) n=3. Scale bar: 25 µm. **(C)** Immunoblotting was conducted to assess the impact of UBE3A depletion on ESR-β expression during oligodendrocyte differentiation. Protein levels of ESR-β in UBE3A-deficient iOPCs and subsequently differentiated iOLs were quantified, normalized against the loading control GAPDH, and expressed relative to the iOPC + siNC control condition (set at 1.0). n= 4. **(D)** The effects of AC-186 (10 µM) and clemastine (2.5 µM) on OPC proliferation (24 hours) and oligodendrocyte differentiation (3 days) were assessed via immunoblotting. Levels of Ki67 and Myelin Associated Glycoprotein (MAG) were measured in UBE3A-deficient iOPCs and iOLs differentiated from them, normalized against GAPDH, and compared to the iOPC + siNC condition. n=4. **(E)** Immunoblotting to examine the necessity of UBE3A for OPC self-renewal and oligodendrocyte differentiation across different developmental stages. Expression of maturation and myelination markers (MBP, MAG, PLP1, CNPase) was quantified following UBE3A depletion timed from iOPCs to pre-iOLs, normalized against TUBB1, and compared to the iOPC + siNC control condition (set at 1.0). n= 4

We further assessed ESR-β functionality during oligodendrocyte differentiation by comparing the effects of AC-186 with clemastine (*49*) - a compound known to promote oligodendrocyte differentiation. While clemastine enhanced differentiation without affecting proliferation in UBE3A-deficient iOPCs, AC-186 showed no impact on differentiation (**Fig. 5D**), suggesting that the impaired ESR-β signaling specifically affects the proliferation stage (iOPC) but not the differentiation into myelinating oligodendrocytes. These findings, aligned with our prior analyses showing a decline in UBE3A levels in mature oligodendrocytes (**Fig. S3B-C**), suggest that UBE3A may play a lesser role in the differentiation process for myelination. To confirm this, we adjusted the timing of UBE3A depletion across different stages of oligodendrocyte development, transitioning from iOPCs to pre-iOL and onto myelin-producing iOL stages. We observed that initiating UBE3A depletion at the iOPC stage significantly impaired differentiation and myelin production, whereas initiating depletion after the onset of differentiation had a lesser impact on these processes (**Fig. 5E**). Additionally, treatment with AC-186 at various stages revealed that its beneficial effects on differentiation were only observed when administered during the iOPC stage, not after differentiation had begun (**Fig. S8D**). These outcomes further delineate the specific stages of oligodendrocyte development affected by UBE3A activity and ESR-β signaling.

### Assessment of *UBE3A* loss-of-function mutation in AS patient-derived OPCs and consequences for proliferation and differentiation for myelination

To rigorously test whether UBE3A loss is sufficient to impair human OPC homeostasis in a patient-relevant setting, we studied iOPCs generated from an AS patient carrying a truncating, loss-of-function *UBE3A* mutation. These AS iOPCs displayed appropriate morphology and expressed canonical lineage markers, supporting normal specification and baseline cellular identity (**Fig. 6A**). Despite this, AS iOPCs exhibited two key disease-associated abnormalities: a marked reduction in proliferative capacity and attenuated ESRβ signaling (**Fig. 6A-B**). Treatment with the selective ESRβ agonist AC-186 robustly restored proliferation in AS iOPCs toward control levels (**Fig. 6C-D**), supporting the conclusion that impaired ESRβ signaling is a functionally relevant downstream consequence of UBE3A deficiency at the OPC stage.

**Figure 6.**
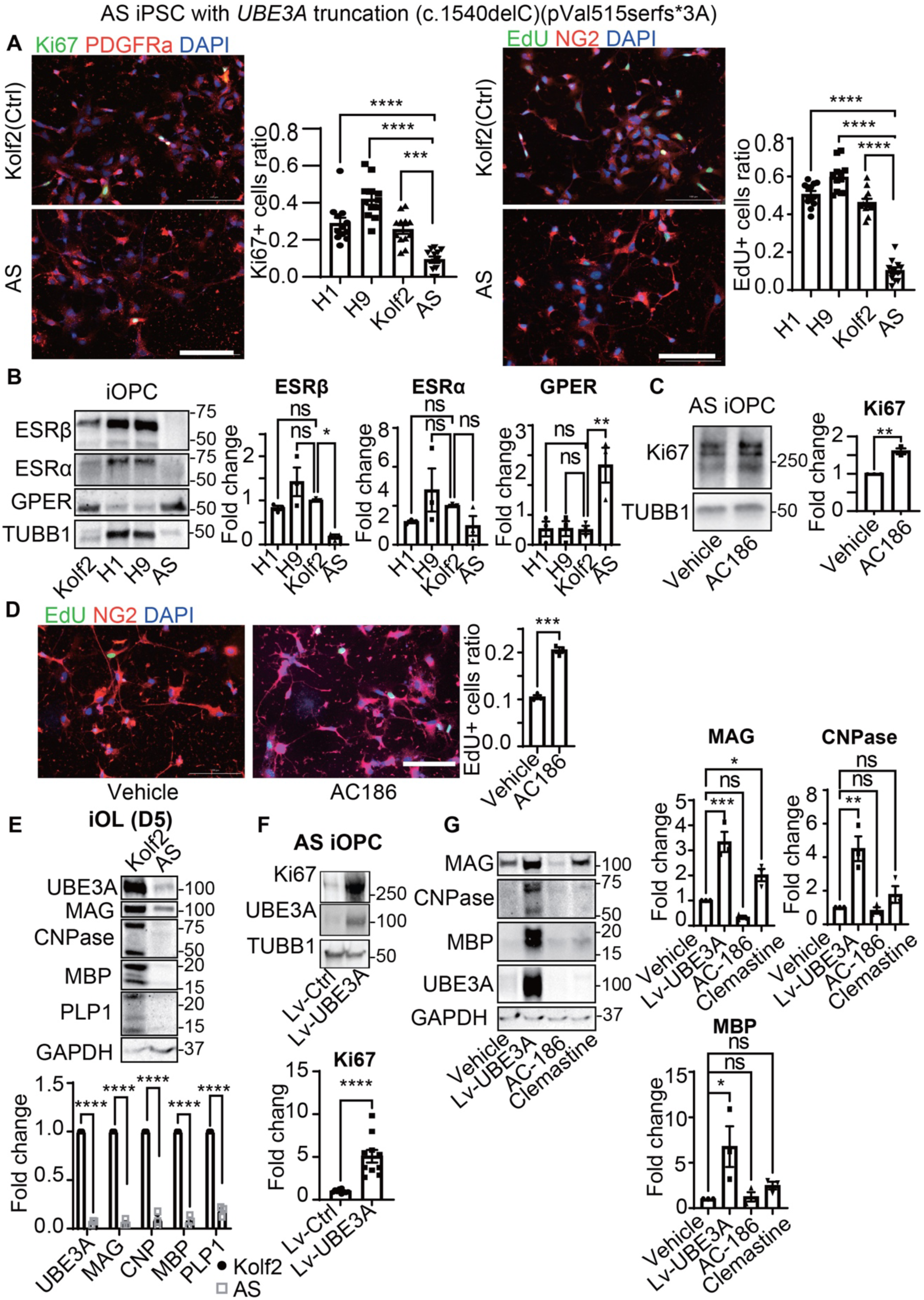
The effect of AS mutation of UBE3A loss-of-function on OPC proliferation and differentiation via ESR-β signaling. **(A)** Imaging validation of iOPCs derived from an AS patient with a UBE3A truncation mutation (AS iOPCs), compared to multiple human pluripotent stem cell lines. OPC markers NG2 and PDGFRα and proliferation markers Ki67 and EdU were immunofluorescently labeled, and OPC proliferation was quantified by the ratio of Ki67+ or EdU+ cells to total OPC cells, in comparison to control iOPCs generated from human embryonic stem cells H1 and H9 and the reference iPSC line Kolf2 lines. n=12 regions of interest from 3 independent experiments. **(B)** Immunoblotting to measure estrogen receptor subtypes ESR-α, ESR-β, and GPER in AS iOPCs, normalizing against TUBB1 and comparing to iOPCs derived from Kolf2, as well as H1 and H9 human embryonic stem cell lines, used as references. Results are presented as fold changes relative to the Kolf2-derived iOPCs (set as 1.0). n=3 experiments. **(C)** Immunofluorescence analysis to characterize the response of AS iOPCs to AC-186 treatment (10 µM, 24 hours) on proliferation, quantified by the ratio of proliferating (EdU+) cells. n=3. **(D)** Immunoblotting to analyze the proliferation effect of AC-186 (10 µM, 24 hours) on AS iOPCs, assessed by quantifying Ki67 expression levels, normalized against β-tubulin (TUBB1), and expressed as fold change relative to the vehicle control condition. n=3. **(E)** Immunoblotting to evaluate the potential of AS iOPCs differentiation into myelinating cells by quantifying the expression of UBE3A and 4 oligodendrocyte markers (MBP, MAG, PLP1, CNPase). Data were normalized against GAPDH and expressed relative to control iOLs derived from the reference iPSC line Kolf2. n=4 experiments. **(F)** Immunoblotting to assess the effect of restoring UBE3A expression on the proliferation of UBE3A-deficient OPCs. In AS iOPCs, UBE3A expression was re-introduced via lentiviral transduction (Lv-UBE3A, 3 days), and Ki67 levels were quantified, normalized to β-tubulin (TUBB1), and compared to a control condition using an empty lentiviral vector (Lv-Ctrl). n=10. **(G)** The effects of restoring UBE3A expression and treatments with AC-186 or clemastine on oligodendrocyte differentiation in AS. AS iOPCs were differentiated into iOLs and treated with lentiviral transduction of UBE3A, AC-186 (10 µM, 72 hours) or clemastine (2.5 µM, 72 hours), with the expression levels of maturation and myelination markers (MAG, CNPase, MBP) quantified. n=3.

To corroborate the pro-proliferative and pro-myelination effects of AC-186 in an independent, genetically defined model, we established primary OPC cultures acutely purified from WT and AS mouse cortex. Consistent with the patient-derived iOPC results, AC-186 selectively rescued the proliferation deficit in AS primary OPCs while exerting minimal effect on WT OPC proliferation (**Fig. S9A**). When these primary OPCs were differentiated in vitro toward myelinating oligodendrocytes, AS cultures showed reduced myelin marker expression, which was again restored by AC-186 to near WT levels, with little to no effect in WT cultures (**Fig. S9B**). Together, these complementary human and mouse primary-cell data strengthen the conclusion that AC-186 preferentially normalizes UBE3A-deficient oligodendroglial phenotypes rather than broadly stimulating proliferation or myelin programs in otherwise normal cells.

We next asked whether ESRβ activation influences later stages of oligodendroglial maturation beyond OPC self-renewal. Upon differentiating AS iOPCs into mature oligodendrocytes, we observed decreased expression of myelination-associated proteins (**Fig. 6E**). To parse stage-specific requirements, we reintroduced UBE3A in AS iOPCs via lentiviral transduction and compared the effects of UBE3A re-expression with AC-186 and with clemastine. UBE3A re-expression robustly improved both OPC proliferation and downstream differentiation into myelinating cells (**Fig. 6F-G**). In contrast, AC-186 primarily corrected the proliferation defect without substantially enhancing differentiation, whereas clemastine increased differentiation into myelinating oligodendrocytes (**Fig. 6G**). These results support a model in which UBE3A regulates multiple stages of oligodendroglial development, while ESRβ agonism acts most prominently at the OPC homeostasis/self-renewal stage.

Collectively, these findings validate a critical requirement for UBE3A in maintaining oligodendroglial homeostasis and establish ESRβ signaling as a therapeutically actionable pathway that selectively restores OPC proliferation and downstream myelin output under UBE3A-deficient conditions. By clarifying the stage specificity of ESRβ activation, improving OPC self-renewal without broadly forcing differentiation, this work highlights a complementary strategy to address oligodendrocyte dysfunction in AS that appears to stem, in large part, from disrupted OPC homeostasis.

### ESR-β activation mitigates oligodendroglial and behavioral deficits resulting from UBE3A depletion in AS mice

After the findings on AS patient-derived iOPCs in response to ESR-β agonist treatment, our investigation then turned to whether AC-186 would remedy the defective OPC homeostasis in vivo and rescue the behavioral abnormality of AS mice. We first examined whether the loss of UBE3A and the consequent reduction in ESRβ function observed *in vitro* could also be detected in the developing brains of AS mice *in vivo*. We evaluated the presence of ESR-β in oligodendrocytes using confocal microscopy to visualize the co-localization of ESR-β immunofluorescence with established markers of oligodendroglia. While the population of oligodendrocyte lineage cells shrank in AS mouse brains (**Fig. 7A**), we observed a significant decrease in ESR-β expression in young AS mouse oligodendrocyte lineage cells (**Fig. 7B**).

**Figure 7.**
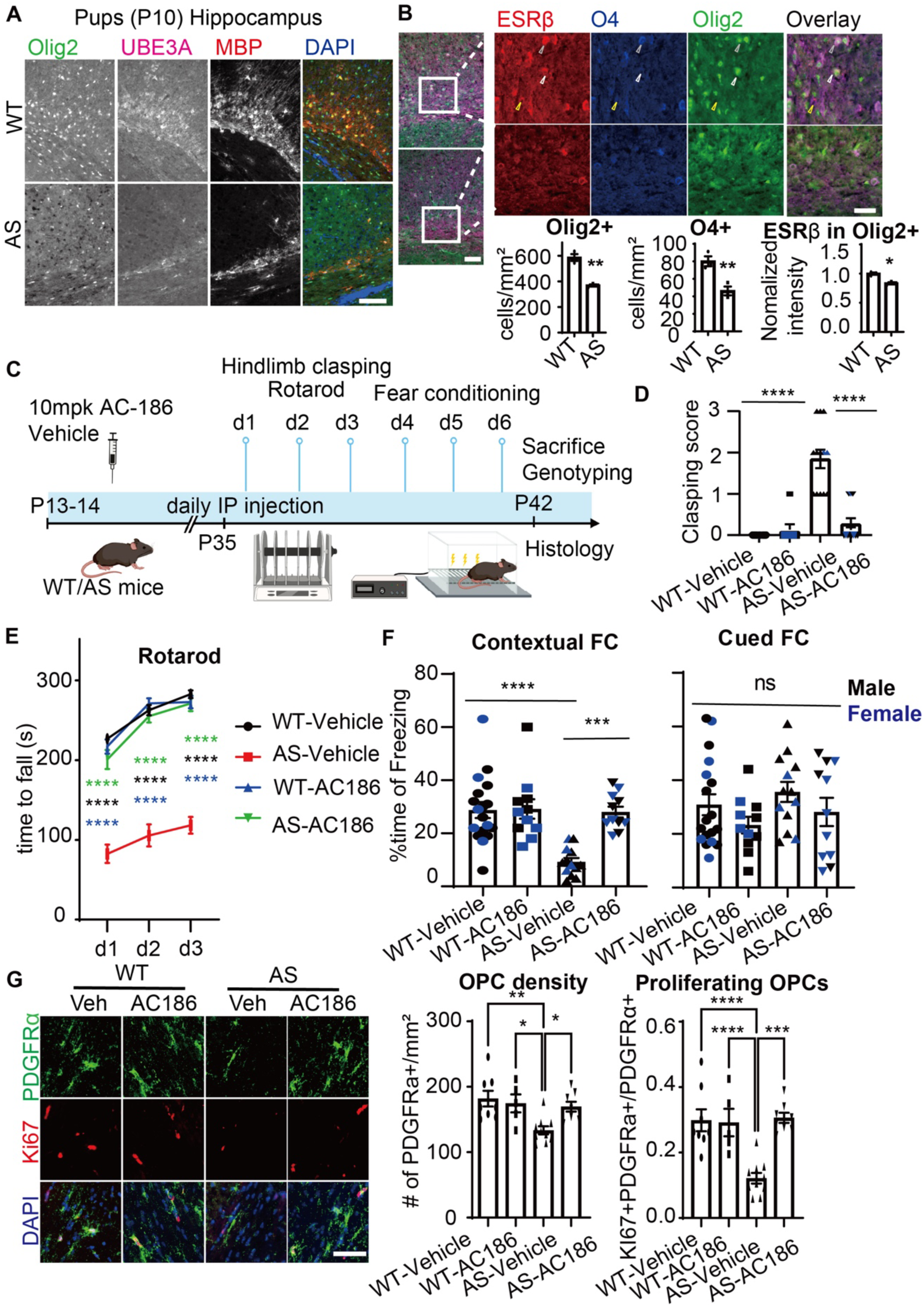
Restoration of oligodendroglial homeostasis and learning behaviors through ESR-β activation in a UBE3A-deficient AS mouse model. **(A)** Brain sections from wildtype and AS mice at early postnatal days (P10) were subjected to immunofluorescence to examine oligodendrocyte populations and UBE3A expression in the hippocampus CA1 region. Olig2 (green) and myelin basic protein (MBP, red) were stained to identify oligodendrocyte lineage cells, co-stained with UBE3A (magenta), and nuclei visualized with DAPI (blue). Scale bar: 100 µm. **(B)** Quantification of oligodendrocyte populations and ESRβ expression. Cell densities for Olig2-positive and O4-positive oligodendrocytes were calculated (left and middle), and ESRβ expression was quantified by fluorescence intensity within Olig2-positive cells. Data from 3 animals per group were analyzed. Scale bars: 100 µm in left panel, 25 µm in right panel. **(C)** Timeline for ESRβ agonist AC-186 efficacy study in AS mice. Juvenile WT or AS mice received daily intraperitoneal injections of ESRβ agonist AC-186 (10 mg per kg of body weight) or a vehicle solution starting at P13-14. Behavioral assessments commenced three weeks later over six days: Hindlimb Clasping (P35; d0), Rotarod (d1-3), and Fear Conditioning (d4-6) tests, commenced. Post-behavioral analysis (P42), mice were sacrificed for immunofluorescence studies of brain sections to quantify oligodendroglial cell density, estrogen receptor expression, and myelination status. Investigative groups were WT with vehicle, WT with AC-186, AS with vehicle, and AS with AC-186. **(D)** Scoring of Hindlimb Clasping. Black symbols, male; blue symbols, females. The number of animals per group was 20 for WT-vehicle, 11 for WT-AC186, 13 for AS-vehicle, and 11 for AS-AC186. **(E)** The motor coordination and learning assessed by the Rotarod test, measuring the time until mice fell or were dislodged from the rotating barrel across all four groups, WT with vehicle (black), WT with AC-186 (blue), AS with vehicle (red) and AS with AC-186 (green), from days 1-3. **(F)** The results of Contextual Fear Conditioning, presented as the percentage of time spent freezing post-tone/shock. **(G)** Examination of OPC population and proliferation in wildtype and AS mouse brains after treatment with vehicle or AC-186; brain sections were immunostained for OPC marker PDGFRα (green), proliferation marker Ki67 (red) and nuclear marker DAPI (blue) to visualize total and proliferating populations of OPCs in the corpus callosum; OPC population quantified as their density, calculated by the number of PDGFRα-positive and DAPI-stained OPCs per square millimeter of corpus callosum area (top bar graph); OPC proliferation is measured as the proportion of dividing OPCs (indicated by Ki67 and PDGFRα dual positivity) relative to the total OPC count (bottom bar graph). n=6-10 animals per group. Statistical significance assessed using Student’s t-test (B), One-way ANOVA (D, F and G), or Two-way ANOVA (E) and subsequent post-hoc tests; * p<0.05, ** p<0.01, *** p<0.001, **** p<0.0001, ns not significant.

We proceeded to explore the restoration of oligodendroglial homeostasis via targeted augmentation of diminished ESR-β signaling as a potential therapeutic approach. The ESR-β agonist AC-186, known for its effective bioavailability in various animal models including rodents and canines (*69–71*), was administered to both male and female juvenile AS mice (**Fig. 7C**). We utilized our established protocols of behavioral assays (*90, 91*) to assess drug efficacy, focusing on phenotypes characteristic of AS mice, such as deficits in contextual fear learning (*5, 73, 92*) and motor learning impairments measured by the accelerating rotarod test (*5, 64, 93, 94*). Additionally, we employed the Hindlimb Clasping test, an observational measure indicative of CNS pathology (*95*). The Hindlimb Clasping test effectively differentiated AS mice from wildtype controls, providing a reliable phenotypic indicator (**Fig. 7D**). Notably, AC-186 treatment significantly alleviated neurological symptoms in AS mice, enhancing their performance to wildtype levels without affecting wildtype mice (**Fig. 7D**). Our three-day Accelerating Rotarod protocol highlighted baseline motor learning impairments in AS mice and revealed significant improvements in AS mice, both male and female, treated with AC-186 compared to those receiving a vehicle solution (**Fig. 7E, S10A**). In the contextual fear conditioning test, juvenile AS mice showed pronounced deficits, which were ameliorated by AC-186 treatment, with no difference between genders (**Fig. 7F**). Conversely, the cued fear conditioning test showed no intergroup differences, suggesting this aspect of fear memory remains unaffected by the syndrome (*5*) (**Fig. 7F**).

After conducting behavioral assessments, we undertook detailed histopathological examinations to evaluate the structural integrity and functional state of oligodendrocyte lineage cells, particularly OPCs and myelin production, alongside the expression of ESR-β in critical brain areas. Our primary analyses centered on OPCs due to their dynamic responsiveness and sensitivity to genetic and environmental stresses in disease conditions (*27*). Prior to the administration of AC-186, we observed a notable reduction in OPC density across various brain regions of AS mice compared to wildtype controls, (**Fig. 7G**). This reduction was quantitatively assessed by counting dividing OPCs, revealing a more pronounced decrease (**Fig. 7G**). Further investigations revealed a decrease in both immature and mature oligodendrocyte lineage cells across gray and white matter regions, marked by specific developmental stage markers, indicating a decline in their populations as well as a deficiency in myelination within AS mouse brains (**Fig. S11A-C, S12A-B**), paralleling reduced *Ube3a* expression in the oligodendrocyte lineage (**Fig. S11D**). These findings confirm a disruption in the homeostatic proliferation of OPCs in AS, impacting both the maintenance of OPC pools and the availability of precursor cells necessary for differentiation into myelinating oligodendrocytes. Our evaluation of the therapeutic effects of the ESR-β agonist AC-186 revealed that while it had no impact on wildtype mice, it notably restored oligodendrocyte populations and myelination in AS mouse brains (**Fig. 7G, S11A-C**). Notably, AC-186 slightly increased ESR-β expression in AS oligodendroglia (**Fig. S11B**), but did not modify UBE3A levels (**Fig. S11C**). Overall, the convergence of our histopathological observations and the enhancements seen in behavioral assays lead us to conclude that the activation of ESR-β could be an effective treatment strategy for disrupted OPC homeostasis.

## DISCUSSION

This study unveils a novel intrinsic mechanism of oligodendroglial homeostasis modulation via UBE3A, a paradigm shifts from the traditionally understood extrinsic influences like polypeptides from neighboring cells. We demonstrate that UBE3A, known for its regulation of cell proliferation (*3*) and links to oncogenesis (*96*) but primarily studied for its roles in neuronal function and pathology, is essential in controlling OPC homeostasis for myelination through a cell-autonomous pathway. Specifically, our results specifically indicate that UBE3A loss disrupts ESRβ signaling in OPCs, crucial for their self-renewal, without impacting the signaling in differentiated oligodendrocytes, thus not directly affecting their differentiation process. By employing iPSC-derived OPCs and oligodendrocytes from AS patients and relevant mouse models, we elucidated that selective ESRβ activation treatment not only corrects these proliferative deficits but also ameliorates related behavioral abnormalities, paving the way for new therapeutic strategies aimed at restoring oligodendroglial homeostasis for brain disorders hallmarked by oligodendrocyte dysfunction.

Oligodendrocyte precursor cells (OPCs), also known as NG2-glia, constitute approximately 5% of total brain cells and are uniformly distributed across both white and gray matter throughout the brain and spinal cord (*24–26*). These cells are dynamic, continuously reorienting their processes and moving through brain parenchyma to actively respond to changes in their environment, similar to microglia (*27*). This dynamic response to neuronal activity enables OPCs to differentiate into myelin-producing oligodendrocytes (*97–101*), an orchestrated process essential for learning and memory functions (*51, 52*). Acute OPC depletion in mouse brains can impair learning behaviors, such as spatial memory (*102*) and motor learning (*103*), in a timeframe much shorter than that required for new myelin formation (*104, 105*), highlighting the crucial role of OPC homeostasis in cognitive development. Although new OPCs can be generated from subventricular zone (SVZ) progenitors (*106*), the maintenance of OPC populations largely relies on local cell interactions, including self-repulsion from cellular contacts (*28*). All OPCs appear capable of entering the cell cycle (*26, 107*), with replacement occurring through the homeostatic behavior of the entire population rather than mobilization of SVZ progenitors or a subset of highly proliferative OPCs. However, this understanding, primarily centered on extrinsic factors like PDGF and its PDGFRα receptor signaling (*30–32*), falls short to explain the altered self-renewal and disrupted homeostasis observed in conditions like chronic demyelination (*34, 35*) or neurofibromatosis (*36*).

Understanding cellular homeostasis is crucial for proper brain function, which relies on the meticulous regulation of various proliferative precursor cells to adapt to rapid and dynamic environmental changes. Our research highlights an OPC-intrinsic mechanism that regulates oligodendroglial homeostasis without impacting oligodendrocyte differentiation. This discovery aligns with live imaging studies in mice (*27*), which showed existing OPCs differentiating into myelinating oligodendrocytes while neighboring OPCs self-renew to replace, thus maintaining the population. This indicates a clear delineation between homeostatic self-renewal and the differentiation program of oligodendrocytes. Significantly, the identification of ESRβ signaling as a regulatory factor opens avenues for therapeutic interventions targeting oligodendrocyte dysfunction and myelin loss. The availability of selective, potent, and safe ESRβ agonists (*72*), already promising in pre-clinical studies for other brain disorders, supports their potential therapeutic application in addressing these neural challenges.

We have found the notion of haploinsufficiency particularly compelling in the context of UBE3A expression and AS. Genomic imprinting of the chromosome region 15q11-q13, leading to the silencing of the paternal *UBE3A* allele in specific brain regions (*5*), has been well-established as the primary mechanism underlying AS. Our findings here suggest that haploinsufficiency - with a single copy of functional *UBE3A* allele failing to achieve a normal phenotypic outcome - also contributes to the disease manifestations of neurodevelopmental delay. Specifically, our research indicates that a moderate reduction in UBE3A expression, as in the situation of loss of one functional *UBE3A* allele, has significant implications for cell types that normally express UBE3A from both parental alleles (*37*), particularly oligodendrocytes (*7*). This understanding is pivotal as it suggests that restoring the function of just one allele might not be sufficient for the optimal functioning of these cells. Recognizing the impact of haploinsufficiency on cellular function expands the scope of potential therapeutic targets and provides a deeper understanding of the pathogenesis of AS, which complements ongoing translational endeavors in unsilencing paternal *UBE3A* in neurons (*12, 13*).

Our findings also underscore the therapeutic potential of selective ESRβ agonists, including AC-186, as a strategy to correct oligodendroglial dysfunction in AS. Given that ESRβ is the pharmacologic target, an additional consideration is whether sex-dependent biology could influence baseline phenotypes or therapeutic response. Importantly, estrogen signaling in the CNS is not solely dictated by circulating gonadal steroids. The brain is a major extra-gonadal source of estrogen, with aromatase expressed in neurons and glia enabling local estradiol production that can act through paracrine and autocrine mechanisms in both sexes (*108*).

Consistent with this, estradiol concentrations within brain regions such as the hippocampus can exceed serum levels, and “brain-derived estrogen” is increasingly viewed as a neuromodulator of synaptic plasticity and cognition rather than simply an endocrine signal that differs by sex (*109, 110*). Moreover, studies in ERβ knockout mice report neurodevelopmental defects -including impaired neuronal migration and survival - in both male and female animals (*111*), supporting a role for ERβ-mediated signaling in CNS development across sexes (*112*). In our datasets, we did not detect an overt sex difference in AS myelination phenotypes; nevertheless, larger sex-stratified cohorts will be important to rigorously test for subtler sex-dependent effects on myelination and on the magnitude or durability of pharmacologic rescue.

We also note several important interpretive considerations. First, OPC proliferation is likely controlled by multiple convergent pathways, and our findings should not be taken to imply that ESRβ signaling is uniquely privileged among all mechanisms regulating OPC homeostasis. Second, because our initial compound screen was conducted at a single concentration, both nonspecific pharmacologic effects and false negatives are plausible - additional pathways may influence proliferation but could be missed without a broader dose-response framework.

Accordingly, we interpret the screen as a hypothesis-generating entry point rather than a comprehensive map of OPC regulators. We prioritized AC-186 for deeper mechanistic and translational follow-up not simply because it produced a strong mean effect in the primary screen, but because its rescue was reproducible across independent culture batches, confirmed in dose-response validation, and supported by convergent evidence from multiple structurally distinct ESRβ agonists and EC50 determination, collectively arguing for an on-target, mechanism-linked effect. Finally, we emphasize that these data do not exclude additional neuronal contributions *in vivo* - ESRβ is expressed in neurons, and UBE3A deficiency alters other estrogen-related nodes (including GPER upregulation also in iNs in our dataset), raising the possibility of receptor crosstalk or compensatory signaling that warrants future targeted investigation.

In conclusion, the importance of this project lies in its potential to directly impact the therapeutic landscape for AS by providing unique insights into the intrinsic role of UBE3A in oligodendrocyte function. By revealing that UBE3A influences OPC dynamics via ESRβ signaling, we provide a pathway that could be therapeutically exploited to ameliorate conditions marked by oligodendrocyte dysfunction. The potential of selective ESRβ agonists to correct OPC proliferative deficits and alleviate associated behavioral impairments offers a promising therapeutic avenue. Our research not only pivots from the classical understanding of OPC regulation by external cues but also provides a foundation for future interventions complementary to ongoing translational endeavors.

## MATERIALS AND METHODS

### Statistical analysis

Sample sizes were determined based on a combination of published effect-size and power estimates for key behavioral readouts (including accelerating rotarod in AS mouse models), prior literature (*64*), and practical considerations (e.g., availability of animals/cultures and experimental batching). Sample sizes for each experiment are reported in the corresponding figure legends.

The normality of data distributions was assessed using the Shapiro-Wilk test, where a p-value less than 0.05 was indicative of a non-normal distribution. The normality of the data distribution was routinely determined by a Shapiro-Wilk normality test (p<0.05 indicating a non-normal distribution). For data confirmed to be normally distributed, pairwise comparisons were conducted using Student’s t-test, while comparisons involving three or more groups were analyzed using One-way or Two-way ANOVA, supplemented by Tukey’s post-hoc test, as specified in the figure legends. In case of non-normally distributed data, we resorted to non-parametric tests, including the Mann-Whitney test or the Kruskal-Wallis test, depending on the data structure. We considered a p-value of less than 0.05 as statistically significant. For data that is not normally distributed, non-parametric alternatives, such as Mann-Whitney or Kruskal-Wallis tests. p<0.05 is considered to be statistically significant.

All graphical representations, including bar graphs and summary plots, display data as means ± standard error of the mean (SEM), derived from a minimum of three independent biological replicates. Significance was reported as * p<0.05, ** p<0.01, *** p<0.001, **** p<0.0001. GraphPad Prism version 9 (GraphPad Software) was utilized for all statistical methods.

## List of Supplementary Materials

Figures. S1 to S12 (embedded in the manuscript)

Table S1 (embedded in the manuscript)

Data files S1 (raw data statistics) and S2 (uncropped images)

## Acknowledgments

We thank our colleagues at Brown University, Dr. Eric Morrow for the helpful discussions and Ms. Gabriela Molica for the primary cultures of neonatal rat oligodendrocyte precursor cells.

## References and Notes

### Funding

This work was supported by the grants from the National Institutes of Health (AG083943; to Y.A.H.), the Foundation for Angelman Syndrome Therapeutics (PD2023-001 to X.Y., and FT2024-001 to Y.A.H.), and the Blackman Family Fund (to Y.A.H.).

### Author contributions

Conceptualization: XY, YAH, YHJ

Methodology: XY, YAH, SM, JM, YHJ

Investigation: XY, YAH

Visualization: XY, YAH

Funding acquisition: XY, YAH

Project administration: YAH, JM

Supervision: YAH

Writing – original draft: XY, YAH

### Competing interests

Y.A.H. is a co-founder of Acre Therapeutics LLC, focusing on the therapeutic research and development of antisense oligonucleotides (ASO) for treatments for tauopathies, including Alzheimer’s. J.M. is a founder of Aingeal, and received funding for research on depression unrelated to the work reported here. The remaining authors declare that they have no conflict of interest.

### Data and materials availability

The complete dataset for this study is included within the main text and Supplementary Materials. For access to raw data, detailed images, and spreadsheets, please refer to the supplementary documentation. Protocols and experimental details are available on request. Material transfers will comply with standard agreements such as the Simple Letter Agreement (SLA) or the Uniform Biological Materials Transfer Agreement (UBMTA) and will exclude reach-through provisions.

---

**Figure S1(related to Fig. 1).**
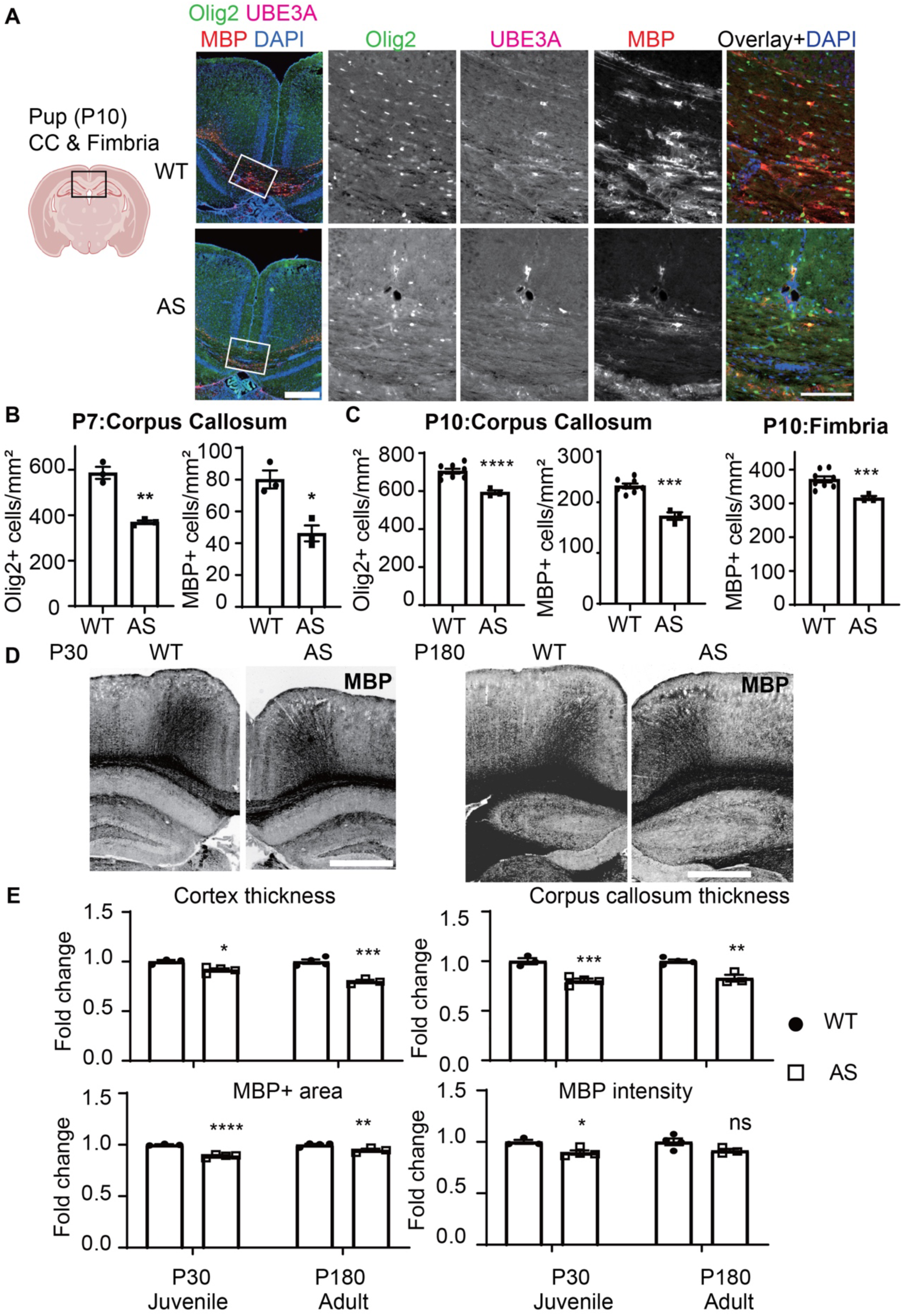
Reduced oligodendrocyte-lineage populations and myelin in AS mouse brains. **(A)** Examination of oligodendrocyte populations and UBE3A expression in the corpus callosum and fimbria of fornix in brain sections from WT and AS pups (P10 shown here). Immunofluorescence staining was performed using Olig2 (green) and myelin basic protein (MBP, red) markers, with concurrent UBE3A (magenta) labeling and DAPI (blue) for nuclear visualization. Scale bars 300µm. **(B-C)** Density metrics of oligodendrocytes within the corpus callosum and fimbria were derived by counting cells positive for Olig2 (left, corpus callosum) or MBP (middle, corpus callosum; right, fimbria of fornix) at postnatal day 7 (P7, **B**) and 10 (P10, **C**). Cell densities were plotted against the assessed area for each marker and data were mean ± SEM. The analysis included 3-8 animals per group. **(D-E)** Examinations of myelination and brain structure in coronal brain sections from juvenile (P30) and adult (P180) WT and AS mice, by visualizing myelin with MBP staining in the gray matter of hippocampus and cortex, as well as the white matter of corpus callosum. Notably, the developing stage of myelination in young pups made MBP detection unreliable, therefore, MBP immunofluorescence was not evaluated in the youngest WT and AS mice (P7-10). n= 3-8 animals per group. Scale bars represent 750µm.

**Figure S2(related to Fig. 1).**
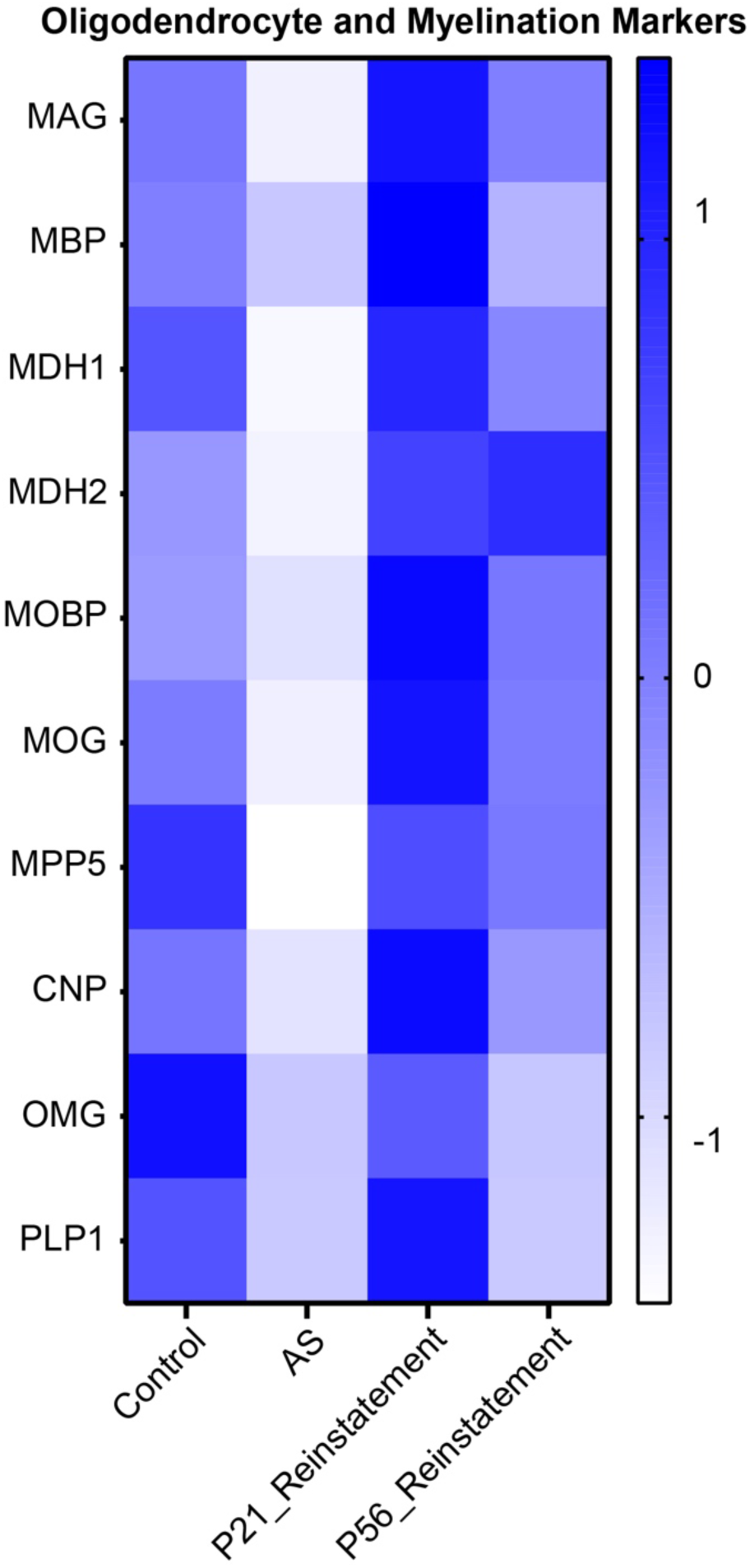
Differential oligodendrocyte and myelination protein expression in a proteome dataset of AS mouse cortical tissues. The heatmap of protein levels indicative of mature oligodendrocytes and myelination in AS using cortical tissue proteome data from an independent AS mouse model that features an inducible reactivation of maternal *Ube3a*, allowing for a stringent analysis of UBE3A function (Pandya *et al.,* 2021). Four groups were included: control mice (WT; Cre^ERT2+^), AS mice (*Ube3a*^Stop/+^; Cre^ERT2-^), and mice with UBE3A reinstatement (*Ube3a*^Stop/+^; Cre^ERT2+^) were injected with tamoxifen at postnatal days P21 (adolescence) or P56 (adulthood) and sacrificed at P84. Cortical tissues from each group were pooled to create a sample-specific spectral library using data-dependent acquisition (DDA) mode, with individual samples subsequently analyzed via data-independent acquisition (DIA) mode.

**Figure S3(related to Fig. 1).**
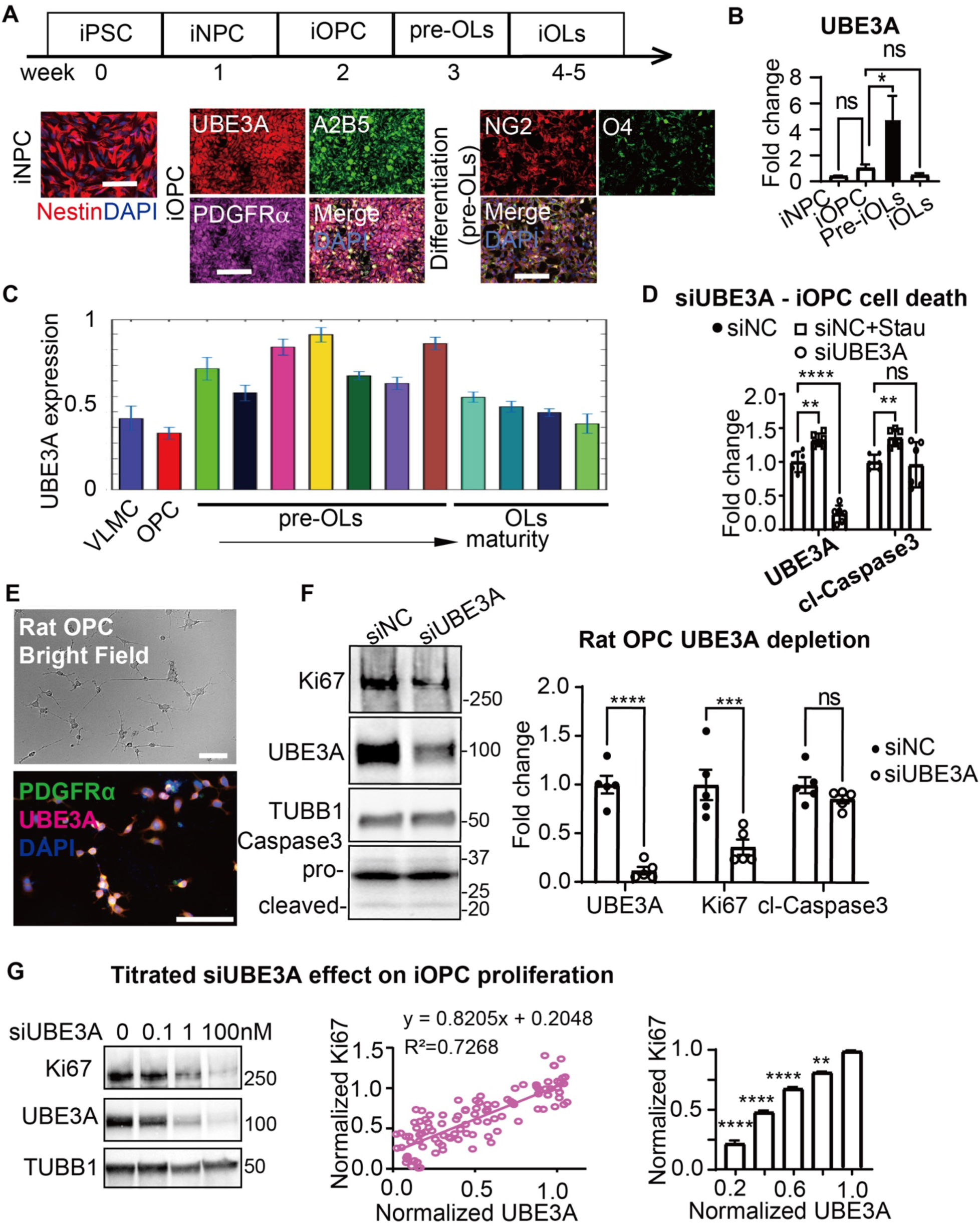
UBE3A expression across developmental stages and its requirement for OPC proliferation. **(A)** Schematic illustrating the process of generating iPSC-derived oligodendrocyte lineage cells using a refined protocol from Vulakh and Yang, 2024(*45*). The human iPSCs employed in this study were the extensively characterized KOLF2.J1 control reference line, sourced from the iNDI(*50*). Initially, iPSCs are converted into neural precursor cells (iNPCs, marked by Nestin) through SMAD inhibition in the first week. These iNPCs are then induced into oligodendrocyte precursor cells (iOPCs, marked by PDGFRα and A2B5) by the second week. Subsequently, iOPCs differentiate into pre-myelinating oligodendrocytes (pre-iOLs, marked by O4) by the third week and finally mature into oligodendrocytes (OLs) by weeks 4-5, ready for myelination assays. Scale bar 100um. **(B)** Assessment of qPCR for UBE3A expression along the oligodendroglial differentiation, at the stages of iNPC, iOPC, pre-iOL and iOL; plotted relative to iOPC (=1.0) after normalizing *UBE3A* mRNA transcript levels to *GAPDH* levels. n=3 experiments. **(C)** Data from single-cell RNA-sequencing of mouse oligodendrocyte lineage cells, as reported by Marques *et al.*, 2016(*46*), showcasing the expression patterns of UBE3A across different stages of oligodendrocyte differentiation in both juvenile and adult central nervous systems. **(D)** Immunoblotting to show no change in cell death of iOPCs undergoing RNA-mediated UBE3A knockdown, using the apoptosis-inducer staurosporine (5 μM for 2h) as a positive control; protein levels of UBE3A and Cl-Caspase were first normalized to β-actin (ACTB) and plotted as relative to the control condition (siNC = 1.0). n=6 experiments. **(E)** Confocal imaging of Bright Field (left) and immunofluorescence (right) to characterize the morphology and functionality of primary OPCs purified from neonatal rat brains and marked by PDGFRα (green) and UBE3A (magenta), scale bar 100um. **(F)** Immunoblotting to evaluate the status of proliferation and cell death in rat primary OPCs treated with siNC (control) or siUBE3A (#1; working for human and rodents) for 2 days, by analyzing the densitometric signals of Ki67 and cleaved form (Cl-Caspase 3) after normalization to loading control tubulin (TUBB1) and plotting as relative to control condition siNC=1.0. The low-activity precursor to Caspase 3 (pro-Caspase 3) could be detected across all conditions. n=5. **(G)** Immunoblotting analysis demonstrating the relationship between UBE3A protein levels and cell proliferation in iOPCs; left panel, by adjusting the dosage of siRNA targeting UBE3A (siUBE3A). Middle Panel: meta-analysis of immunoblotting data for UBE3A and the proliferation marker Ki67 in iOPCs treated with varying doses of siUBE3A. Intensity values were normalized to a loading control and compared against the normalized control (siNC), set as 1.0. A significant positive correlation between UBE3A and Ki67 levels is depicted. Right panel: data presented as a bar graph, categorizing groups based on the degree of UBE3A reduction achieved by RNA-mediated knockdown. Groups exhibiting less than 80% of the mean UBE3A level relative to siNC demonstrated notably lower Ki67 expression compared to the group with the smallest reduction (80-100% of siNC mean). Sample sizes were n=30 for <20% reduction, 20 for 20-40% reduction, 21 for 40-60% reduction, 11 for 60-80% reduction, and 28 for 80-100% reduction, across multiple experiments.

**Figure S4 (related to Fig. 2).**
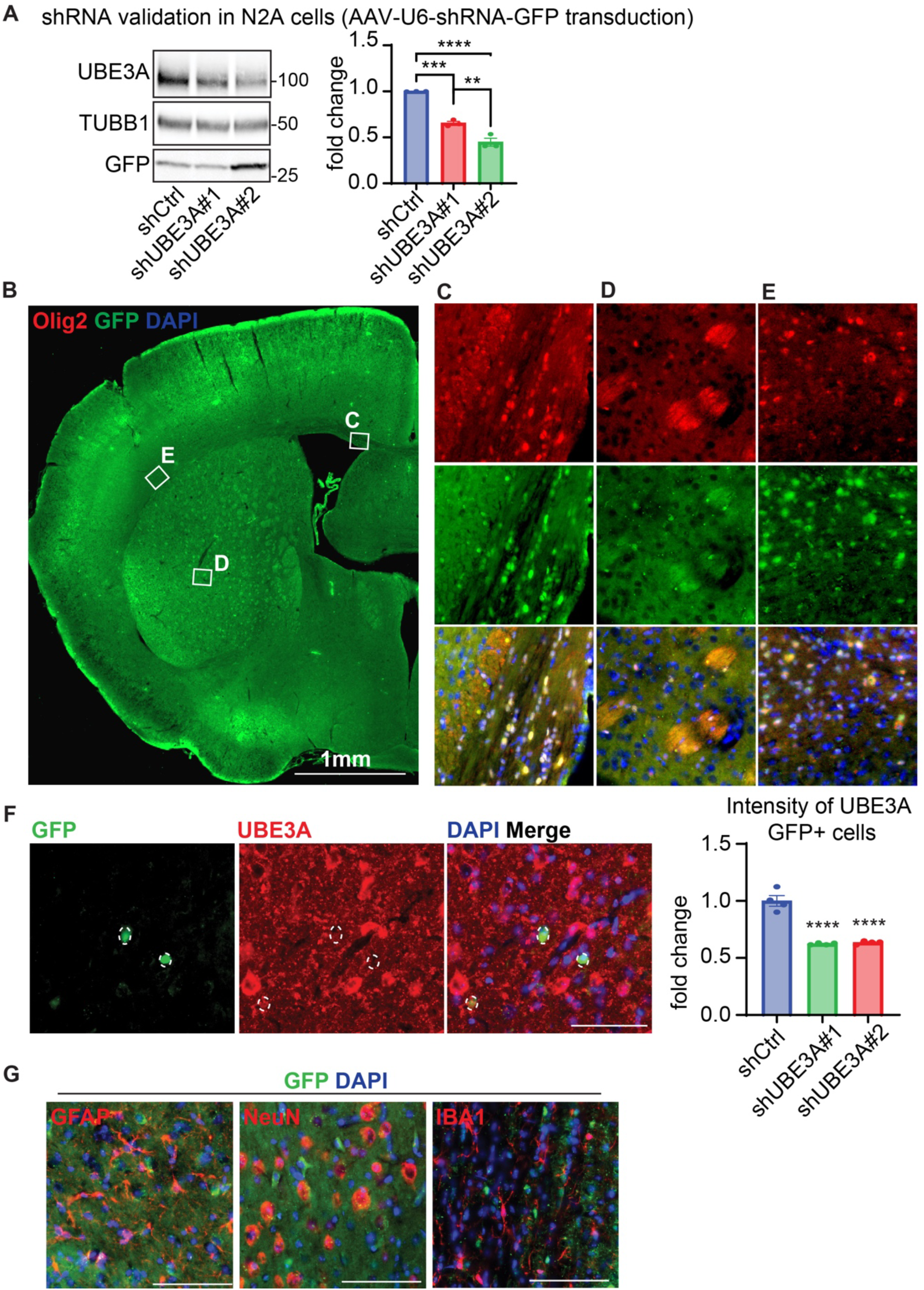
Validation of oligodendrocyte-lineage targeting and *in vivo Ube3*a knockdown by AAV-Olig1-shRNA-GFP. **(A)** Immunoblot validation of shRNA-mediated *Ube3a* knockdown in mouse N2A cells following PEI-based transfection with AAV-U6–shRNA-GFP constructs expressing shCtrl, shUBE3A#1, or shUBE3A#2. UBE3A was quantified and normalized to TUBB1, and values were normalized to shCtrl. *n* = 3 independent experiments. **(B)** Low-magnification overview of a P42 brain hemisphere following P0 intracerebroventricular injection of AAV-Olig1–shRNA-GFP, showing widespread GFP signal. Boxes indicate regions enlarged in panels (C–E). Scale bar, 1 mm. **(C–E)** Higher-magnification views of boxed regions in (B) showing AAV transduction (GFP, green) and oligodendrocyte-lineage cells (Olig2, red), with DAPI nuclear counterstain (blue), illustrating preferential GFP labeling within the oligodendrocyte lineage across distinct brain regions; corpus callosum (**C**), external capsule (**D**), and Striatum (**E**). Scale bar, 200 µm. **(F)** In vivo validation of UBE3A knockdown efficiency by immunofluorescence quantification of UBE3A intensity within GFP+ transduced cells in the hippocampus. UBE3A (red) and GFP (green) were visualized with DAPI nuclear counterstain (blue); UBE3A fluorescence intensity in GFP+ cells was normalized to shCtrl. *n* = 4 animals per group. Scale bar, 200 µm. **(G)** Assessment of cell-type specificity of AAV-Olig1–shRNA-GFP expression by co-immunostaining for GFP with markers of astrocytes (GFAP), neurons (NeuN), and microglia (IBA1), showing minimal GFP overlap with these non-target cell types. Nuclei were counterstained with DAPI (blue). Scale bar, 200 µm.

**Figure S5 (related to Fig. 3).**
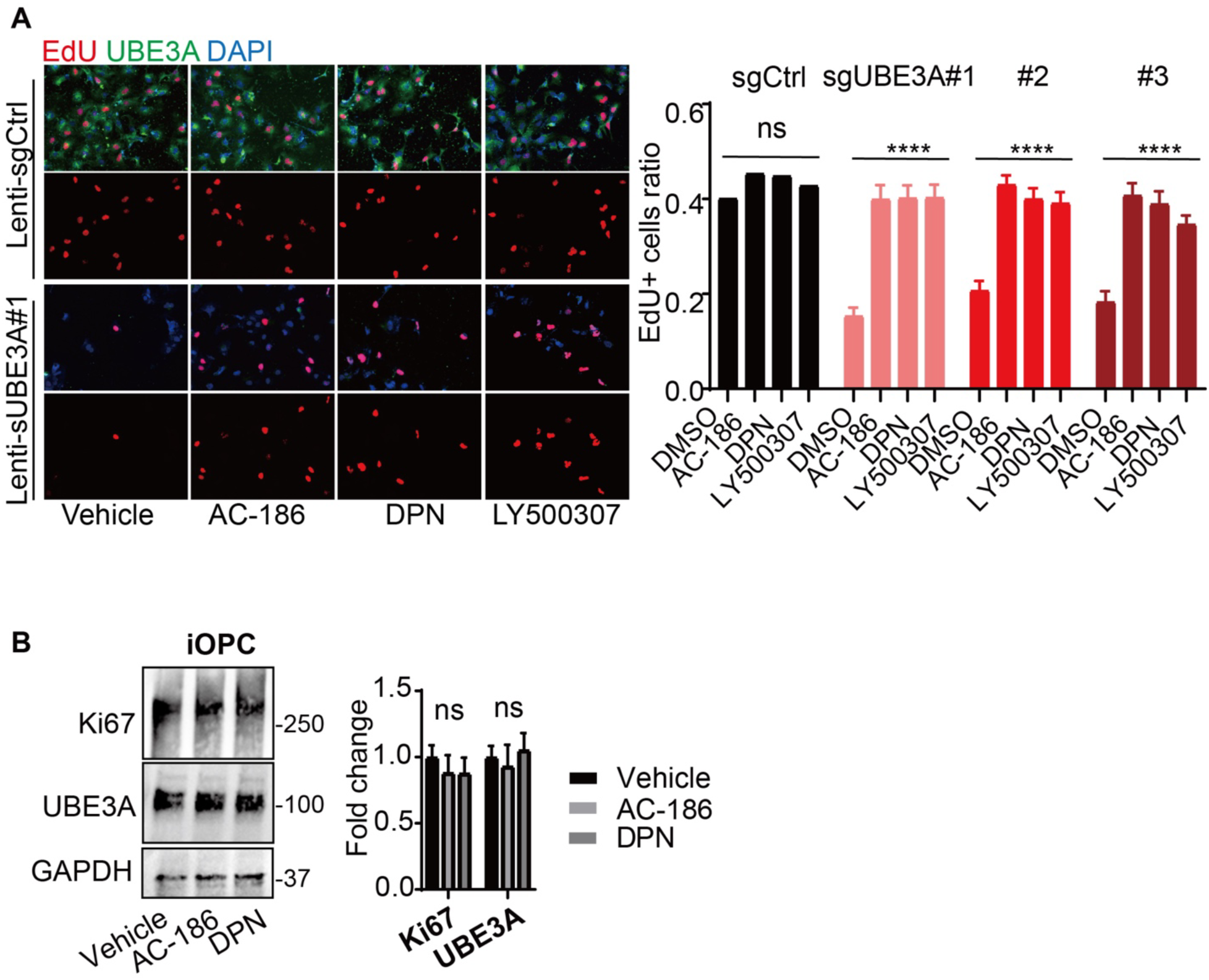
Assessment of UBE3A and ESRβ-mediated OPC proliferation and their pharmacological modulations. **(A)** Immunofluorescent staining to assay the OPC proliferation altered by UBE3A depletion followed by ESRβ stimulation, measuring the EdU (red) nuclear incorporation in iOPCs expressing Cas9 and a sgRNA targeting *UBE3A* via lentiviral transduction (Lenti-sgUBE3A#1-3) or a control sgRNA (Lenti-sgCtrl) and treated with either the vehicle control (DMSO), AC-186, DPN, or LY500307 commenced at a standard concentration of 10 µM, and then labeled with DAPI (blue) and UBE3A (green); data were plotted by the percentage of cells (DAPI) positive for EdU. n= 4 experiments. **(B)** Baseline effects of AC-186 on OPC proliferation analyzed by Ki67 expression in untreated iOPCs exposed to vehicle or drugs (AC-186 or DPN; 10 µM, 24 hours), with densitometric signals of immunoblotting normalized to GAPDH. No significant changes were observed by One-way ANOVA, complemented by post-hoc tests. n=7.

**Figure S6 (related to Fig. 3).**
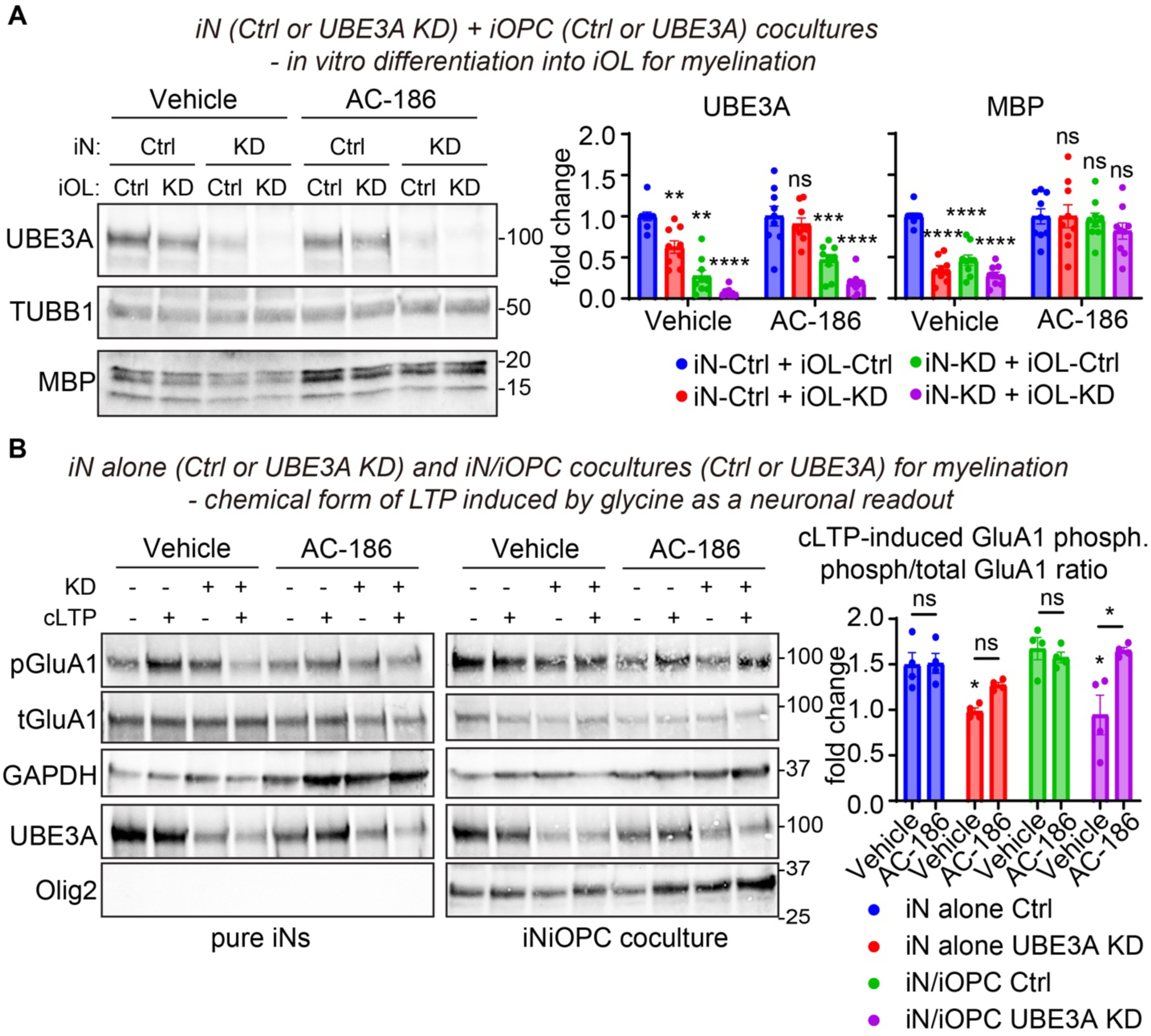
iN–iOPC co-culture resolves cell-intrinsic versus interaction-dependent effects of UBE3A loss and AC-186 on myelination and neuronal plasticity readouts. **(A)** Immunoblot assessment of myelination output in a human iPSC-based co-culture system combining induced neurons (iN; Ctrl or UBE3A KD) with iPSC-derived oligodendrocyte lineage cells (iOPC/iOL; Ctrl or UBE3A KD) and differentiated in vitro to generate myelinating oligodendrocytes. All four combinations were treated with vehicle or AC-186, and myelin output was quantified by MBP expression. UBE3A and MBP were quantified and normalized to TUBB1, with values plotted relative to the vehicle-treated iN-Ctrl + iOL-Ctrl condition (set to 1.0). *n* = 9 cultures from 3 independent experiments. **(B)** Immunoblot assessment of neuronal synaptic plasticity signaling in iN monocultures (Ctrl or UBE3A KD) and matched iN/iOPC co-cultures (Ctrl or UBE3A KD) treated with vehicle or AC-186, followed by chemical LTP induction (glycine). Plasticity signaling was quantified as the pGluA1/total GluA1 ratio and plotted relative to the corresponding control condition (as indicated). GAPDH served as loading control. UBE3A knockdown was verified by UBE3A immunoblotting, and Olig2 was included to confirm the presence of oligodendrocyte-lineage cells in co-culture conditions. *n* = 4 independent experiments.

**Figure S7(related to Fig. 4).**
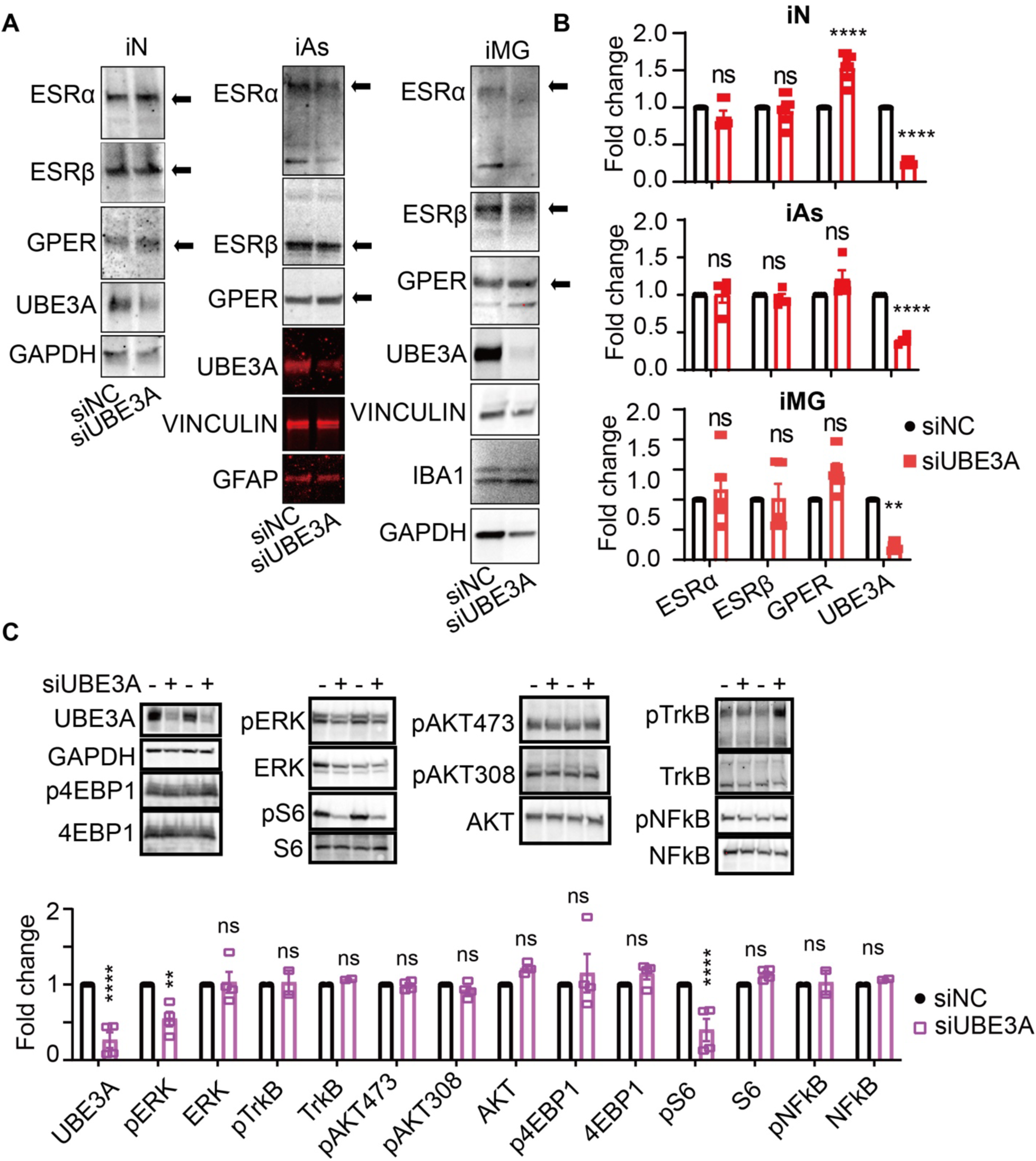
Impact of UBE3A depletion on estrogen receptor (ESR) signaling and downstream pathway activation in neural cells. **(A-B)** Immunoblotting assessed how UBE3A depletion impacts ESR expression in neurons, astrocytes and microglia. Protein levels of ESR-α, ESR-β, GPER, and UBE3A were quantified in human iPSC-derived neurons (iN), astrocytes (iAs) and microglia (iMG) after siNC or siUBE3A treatment, normalized against GAPDH, and compared to siNC (value set at 1.0). n=4-5 replicate experiments; statistical significance was determined by Two-way ANOVA with subsequent post-hoc tests; ** p<0.01, **** p<0.0001, ns not significant. **(C)** Determination of how UBE3A depletion impacts downstream signaling of ESRs, immunoblotting was used to measure the activation of several key signaling pathways in iOPCs treated with siNC or siUBE3A. Pathway activation was assessed by calculating the ratio of phosphorylated to total protein levels for key mediators such as 4EBP1, ERK, S6, TrkB, and NFkB. The efficiency of UBE3A knockdown was verified, and changes in activation status were normalized to GAPDH and compared to siNC (baseline set at 1.0). Four replicates were analyzed for each condition, with statistical significance assessed using Two-way ANOVA and subsequent post-hoc tests; ** p<0.01, **** p<0.0001.

**Figure S8 (related to Fig. 5).**
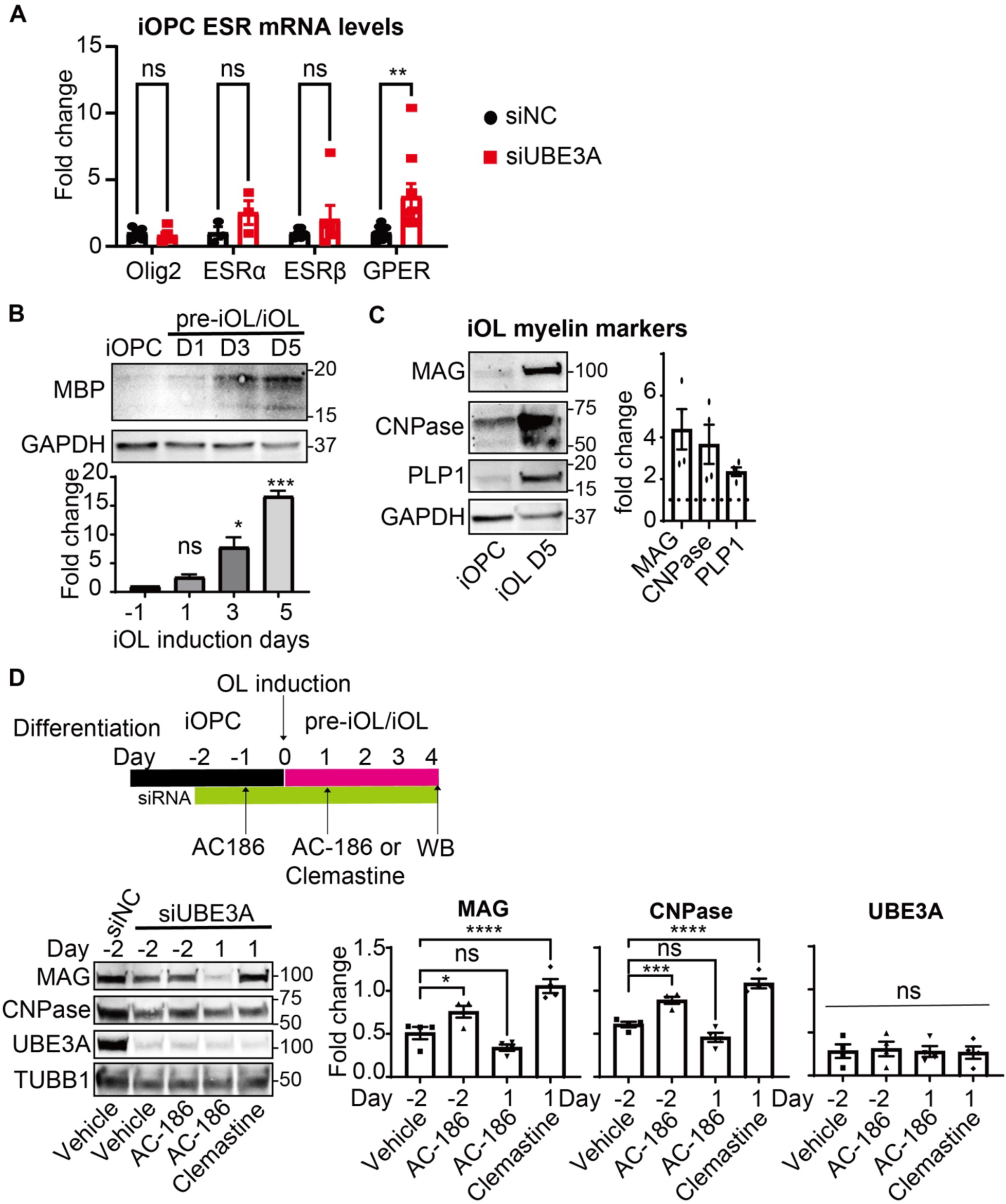
Differential impacts from UBE3A via ESR-β signaling on OPC proliferation and oligodendrocyte differentiation. **(A)** Assessments of qPCR to evaluate the transcriptional control of estrogen receptors in OPCs after UBE3A depletion, by measuring mRNA transcripts of ESR-α, ESR-β, GPER and Olig2 as a negative control in iOPCs treated with siNC or siUBE3A. n=4-6 experiments. **(B-C)** Immunoblotting assessed oligodendrocyte differentiation in iOPCs, following the protocol outlined in Figure S1A. Protein expression levels of myelination markers myelin basic protein (MBP), myelin associated glycoprotein (MAG), 2’,3’-cyclic nucleotide 3’ phosphodiesterase (CNPase), and proteolipid protein 1 (PLP1) were measured. Results were normalized to GAPDH and are presented as fold change relative to the untreated iOPC condition, set as 1.0. n=4. **(D)** Immunoblot analysis of stage-dependent effects of AC-186 (10 µM) and clemastine (2.5 µM) during oligodendrocyte differentiation under sustained UBE3A depletion. iOPCs were transfected with siRNA against UBE3A (or non-targeting siRNA) beginning 2 days before differentiation induction (Day −2) to ensure knockdown at the OPC stage, with repeated siRNA treatments maintained through the transition to pre-iOL and iOL stages. AC-186 was initiated at Day −1 (OPC stage) as indicated, whereas AC-186 or clemastine were alternatively initiated at Day +1 (after differentiation induction; early pre-iOL stage) to compare timing-dependent effects. MAG and CNPase were quantified, normalized to TUBB1 (β-tubulin), and expressed as fold change relative to the siNC control condition (set to 1.0). n=4.

**Figure S9 (related to Fig. 6).**
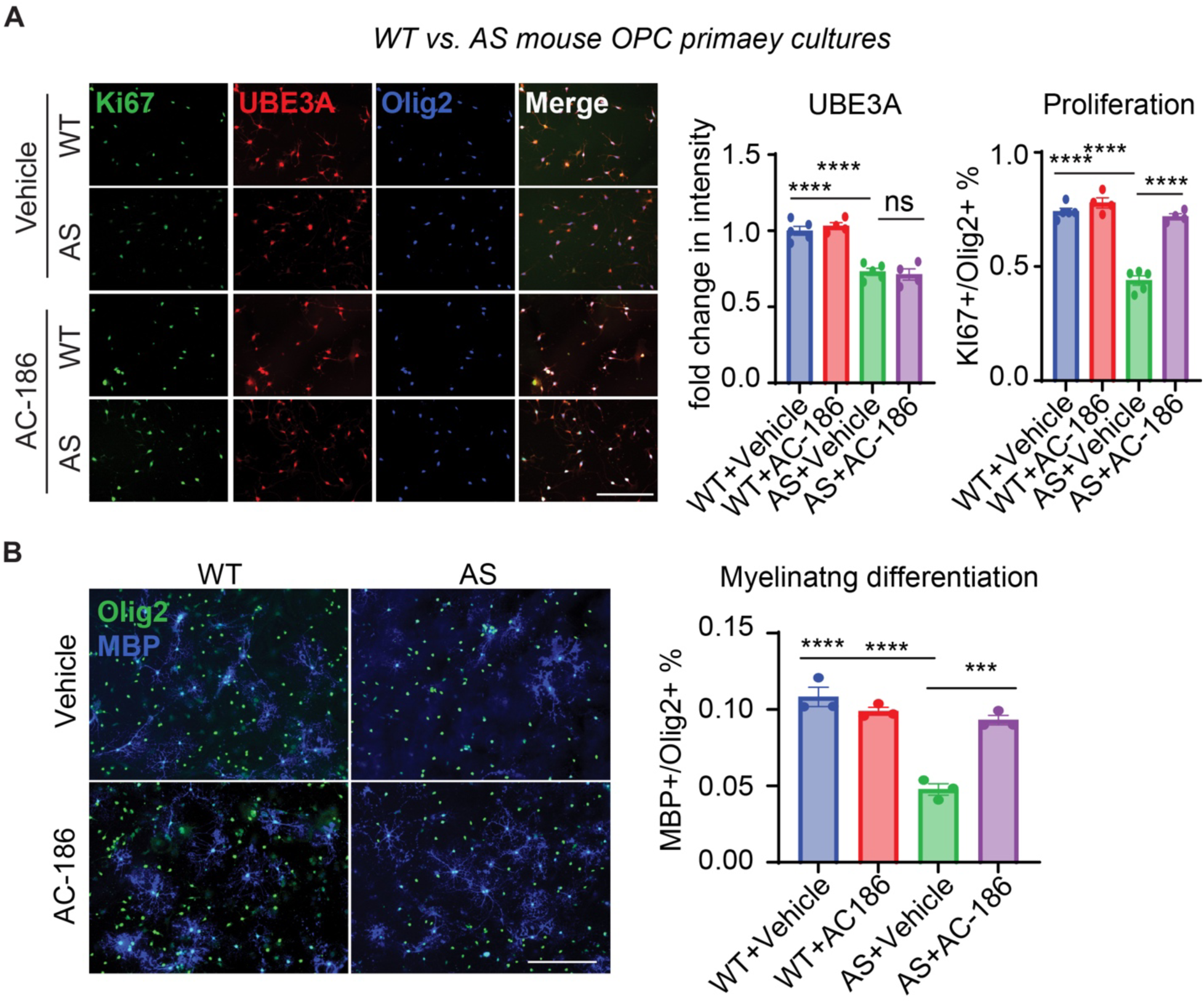
AC-186 restores proliferation and myelinating differentiation in primary OPC cultures from AS mouse cortex. **(A)** Primary OPCs purified from WT or AS mouse cortex were cultured and treated with vehicle or AC-186, then assessed for proliferation by immunofluorescence staining of Ki67 (green) with concurrent labeling of UBE3A (red) and the oligodendrocyte-lineage marker Olig2 (blue). Quantification shown as UBE3A fluorescence intensity and the percentage of Ki67+ cells among Olig2+ cells for each condition. **(B)** Primary WT and AS OPC cultures were differentiated in vitro toward myelinating oligodendrocytes in the presence of vehicle or AC-186 and stained for Olig2 (green) and the myelin marker MBP (blue). Quantification shown as the percentage of MBP+ cells among Olig2+ cells as a measure of myelinating differentiation.

**Figure S10(related to Fig. 7).**
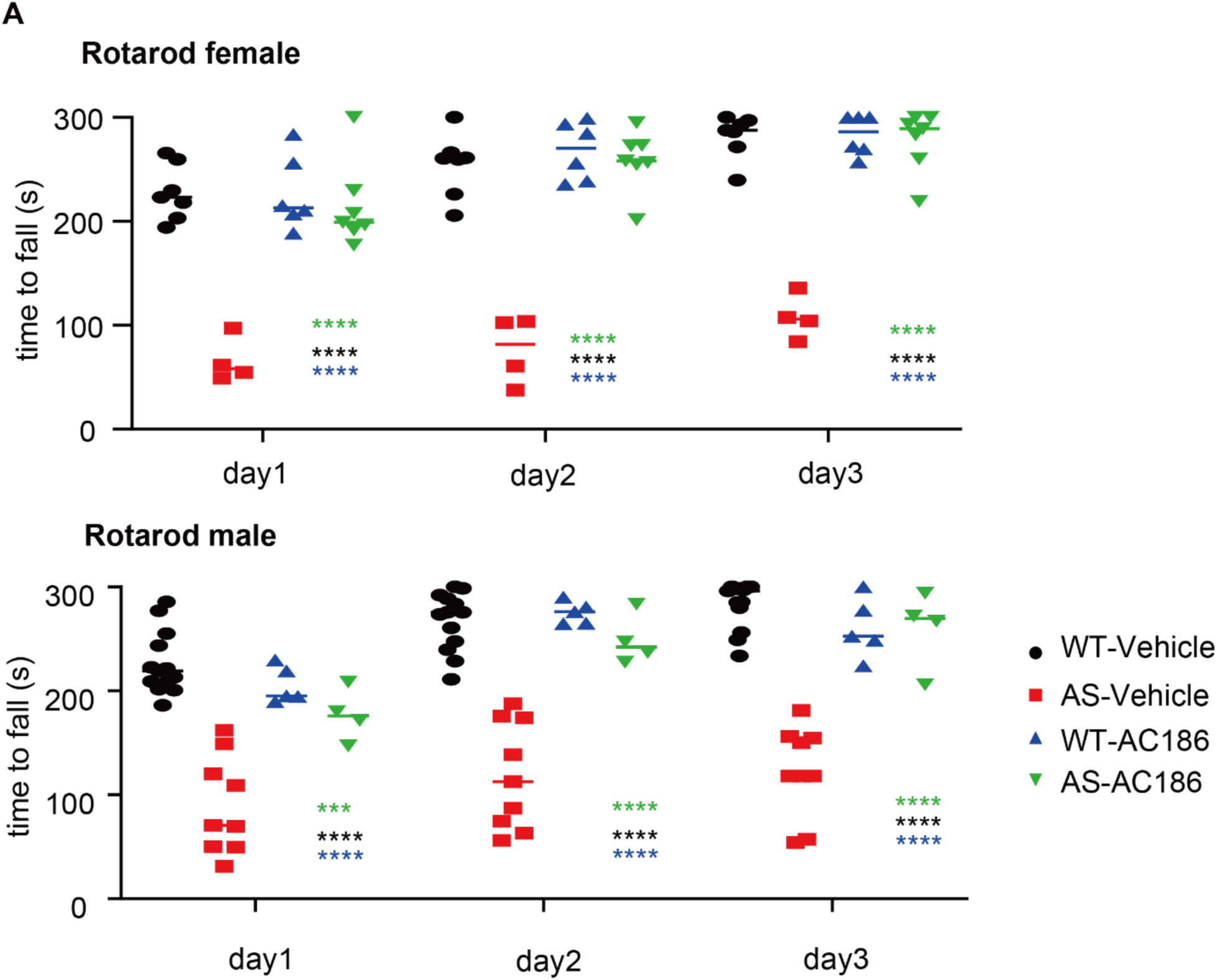
**(A)** Examination of the gender effect on ESRβ response in behavioral assays, by further analyzing the Rotarod data with grouping of female (top) and male (bottom) mice. For female mice, n=7 for WT-vehicle, 6 for WT-AC-86, 4 for AS-vehicle, and 7 for AS-AC186. For male mice, n=13 for WT-vehicle, 5 for WT-AC-86, 9 for AS-vehicle, and 4 for AS-AC186. Statistical significance was determined by Two-way ANOVA with post-hoc tests; *** p<0.001 or **** p<0.0001 for all the comparisons vs. AS-AC186.

**Figure S11(related to Fig. 7).**
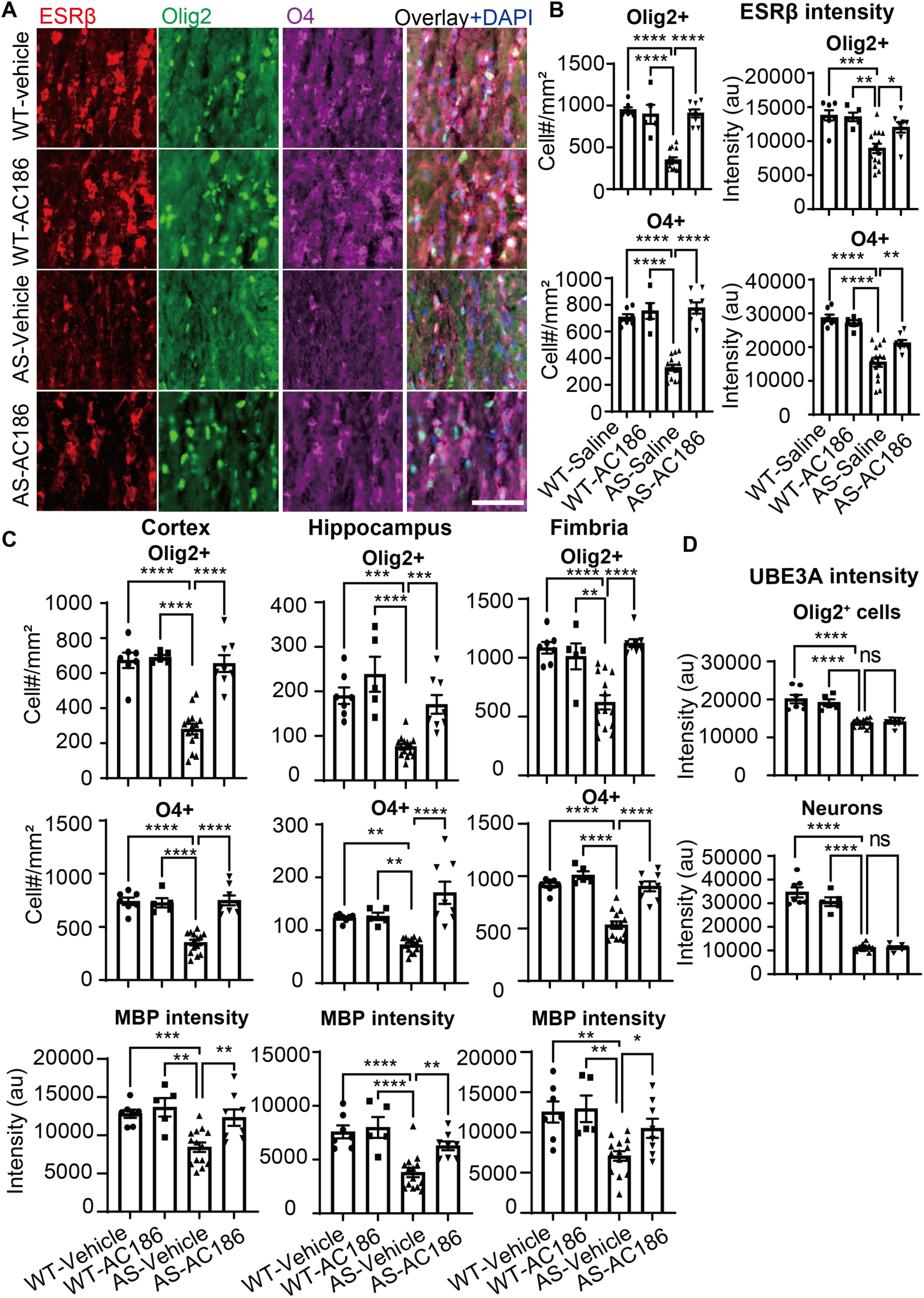
Improved oligodendroglial homeostasis and myelination in treatment-responsive AS mice. **(A)** Immunofluorescence for analysis of oligodendroglia population and ESRβ expression in WT and AS mice treated with vehicle control or AC-186, by imaging the corpus callosum stained with Olig2 (green), O4 (magenta), ESRβ (red) and DAPI (blue). Scale bar: 30 μm. **(B)** Calculations of Olig2-positive and O4-positive oligodendroglial cell densities and their ESRβ expression in the corpus callosum among four groups: WT with vehicle, WT with AC-186, AS with vehicle, and AS with AC-186. n=5-14 animals per condition. **(C)** Comparative assessment of oligodendroglial populations and myelination across brain regions of motor cortex (left column), hippocampus (middle column) and fimbria of fornix (right column), by quantifying the cell densities of Olig2-positive (top row) and O4-positive (middle row) cells as well as the MBP immunofluorescence signal intensity (bottom row). n=5-14 per condition. **(D)** Evaluation of UBE3A expression in oligodendrocyte lineage cells and neuron-like cells, based on morphology and lack of oligodendroglial markers Olig2 and MBP. n=5-14 per condition. One-way ANOVA with post-hoc tests was used to determine statistical significance; * p<0.05, ** p<0.005, *** p<0.001, **** p<0.0001, ns not significant.

**Figure S12 (related to Fig. 7 and S11).**
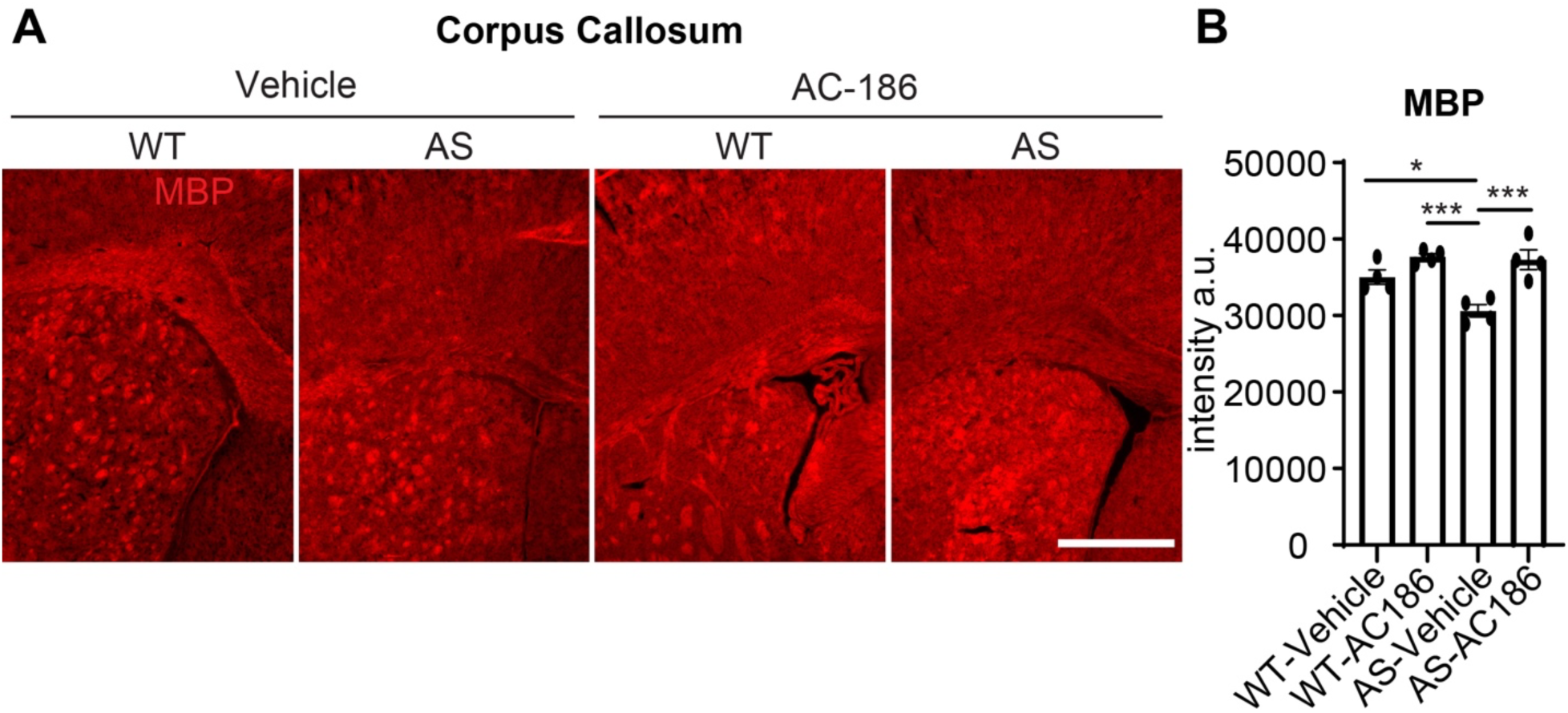
AC-186 improves myelin expression in young adult AS mouse brain (corpus callosum region) **(A)** Representative immunohistochemistry images of MBP staining (red) in the corpus callosum from young adult WT and AS mice treated with vehicle or AC-186. Scale bar: 300 μm. **(B)** Quantification of MBP fluorescence intensity in the corpus callosum for the indicated groups (WT + vehicle, WT + AC-186, AS + vehicle, AS + AC-186). n=4 animals per condition. Each dot represents one animal; bars show mean ± SEM. One-way ANOVA with post-hoc tests was used to determine statistical significance; * p<0.05, *** p<0.001.

**Table S1 (related to Figure 1).**
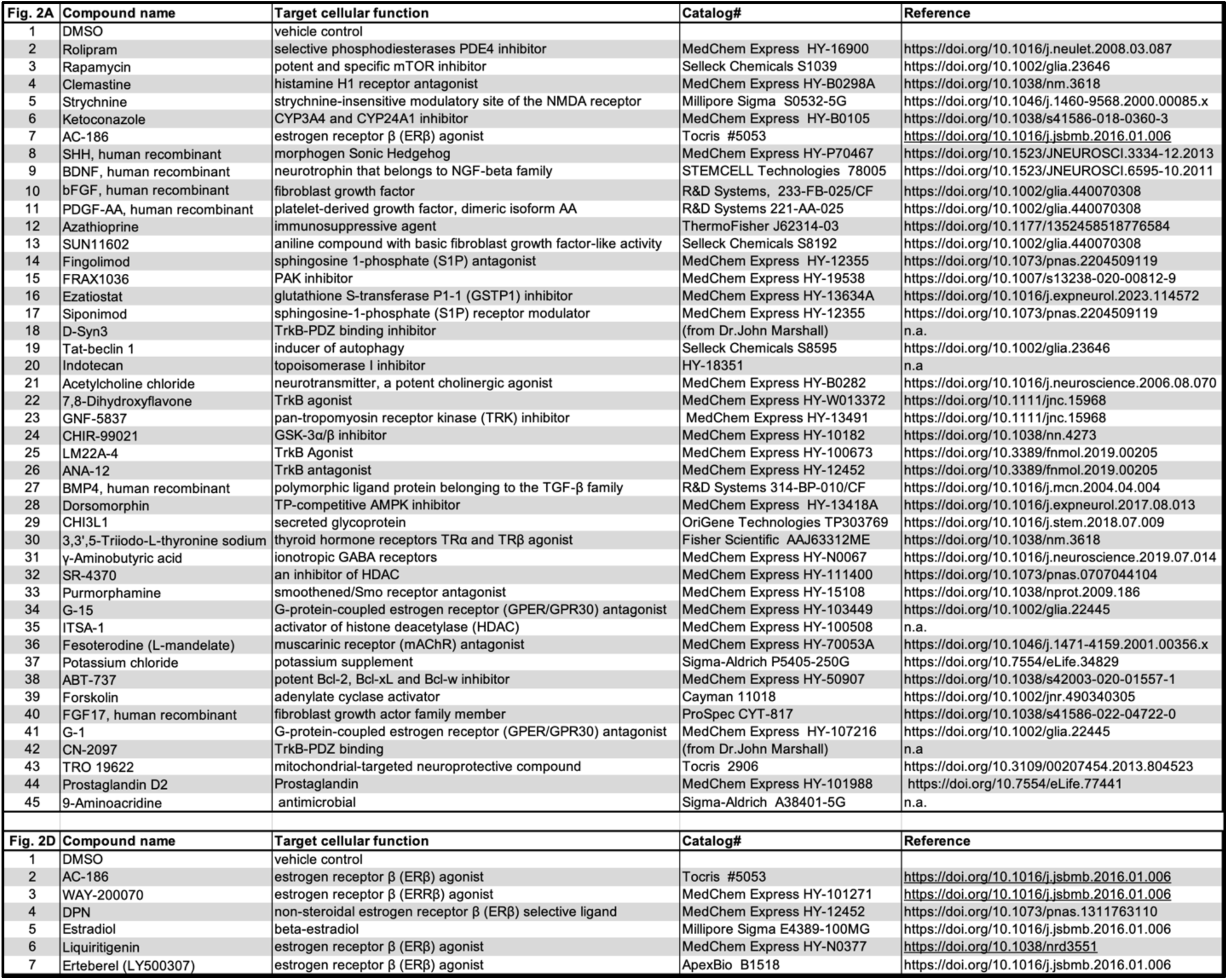
The list of compounds selected for the screen and validation.

